# Molecular mechanism underlying SNARE-mediated membrane fusion enlightened by all-atom molecular dynamics simulations

**DOI:** 10.1101/2023.11.21.568197

**Authors:** Josep Rizo, Levent Sari, Klaudia Jaczynska, Christian Rosenmund, Milo M. Lin

## Abstract

The SNARE proteins syntaxin-1, SNAP-25 and synaptobrevin mediate neurotransmitter release by forming tight SNARE complexes that fuse synaptic vesicles with the plasma membranes in microseconds. Membrane fusion is generally explained by the action of proteins on overall membrane properties such as curvature, elastic modulus and tension, and a widespread model envisions that the SNARE motifs, juxtamembrane linkers and C-terminal transmembrane regions of synaptobrevin and syntaxin-1 form continuous helices that act mechanically as semi-rigid rods, squeezing the membranes together as they assemble (‘zipper’) from the N- to the C-termini. However, the mechanism underlying fast SNARE-induced membrane fusion remains unknown. We have used all-atom molecular dynamics simulations to investigate this mechanism. Our results need to be interpreted with caution because of the limited number and length of the simulations, but they suggest a model of membrane fusion that has a natural physicochemical basis, emphasizes local molecular events over general membrane properties, and explains extensive experimental data. In this model, the central event that initiates fast (microsecond scale) membrane fusion occurs when the SNARE helices zipper into the juxtamembrane linkers which, together with the adjacent transmembrane regions, promote encounters of acyl chains from both bilayers at the polar interface. The resulting hydrophobic nucleus rapidly expands into stalk-like structures that gradually progress to form a fusion pore, aided by the SNARE transmembrane regions and without clearly discernible intermediates. The propensity of polyunsaturated lipids to participate in encounters that initiate fusion suggests that these lipids may be important for the high speed of neurotransmitter release.

---

The release of neurotransmitters by Ca^2+^-evoked synaptic vesicle exocytosis constitutes the primary means of communication between neurons. This exquisitely regulated process involves tethering of synaptic vesicles to the presynaptic plasma membrane, priming of the vesicles to a release-ready state(s) and fast fusion of the vesicles with the plasma membrane upon Ca^2+^ influx into the presynaptic terminal (1). These steps are controlled by a complex protein machinery that has been extensively studied (2, 3), allowing reconstitution of basic features of synaptic vesicle fusion with purified components (4–6) and definition of their functions. The SNAP receptors (SNAREs) synaptobrevin, syntaxin-1 and SNAP-25 play a central role in membrane fusion by forming tight four-helix bundles called SNARE complexes that bring the membranes together (7–10). N-ethylmaleimide sensitive factor (NSF) and soluble NSF attachment proteins (SNAPs) disassemble SNARE complexes to recycle the SNAREs (7, 11). Munc18-1 and Munc13-1 organize SNARE complex formation by an NSF-SNAP-resistant mechanism (4, 12) that involves binding of Munc18-1 to a ‘closed’ conformation of syntaxin-1 (13, 14) and to synaptobrevin, thus templating SNARE assembly (15–17) while Munc13-1 opens syntaxin-1 (18) and bridges the vesicle and plasma membranes (5, 19, 20). The resulting partially assembled SNARE complex binds to synaptotagmin-1 (21) and to complexin (22), forming a spring-loaded primed state (23) that is ready to trigger fast membrane fusion when Ca^2+^ binds to synaptotagmin-1 (24).

While this overall mechanism of neurotransmitter release is well established and it is clear that the neuronal SNAREs alone can induce fusion of reconstituted proteoliposomes (25, 26), a fundamental question remains unanswered: how do the SNAREs induce membrane fusion? Moreover, synaptic vesicle fusion likely occurs in a few microseconds, as the delay between presynaptic Ca^2+^ influx and postsynaptic currents in rat cerebellar synapses is 60 μs (27) and multiple events occur within this time frame. In contrast, cryo-electron microscopy (cryo-EM) images of SNARE-mediated liposome fusion reactions showed that the SNAREs induce extended bilayer-bilayer interfaces that fuse in seconds or minutes (26). It is this unclear how the release machinery induces fusion in the microsecond time scale.

The SNARE four-helix bundle is formed by sequences called SNARE motifs (9, 10). SNAP-25 contains two SNARE motifs, whereas synaptobrevin and syntaxin-1 each contain one SNARE motif that is followed by a short juxtamembrane (jxt) linker and a C-terminal transmembrane (TM) region anchored at the synaptic vesicle or plasma membrane, respectively (Fig. 1*A*). X-ray crystallography showed that the SNARE motif, jxt linker and TM regions of synaptobrevin and syntaxin-1 fusion form continuous α-helices in the cis-SNARE complexes that are expected to be formed after membrane fusion (28) (Fig. 1*B*), and a widespread textbook model envisions that these helices act as relatively stiff rods that exert mechanical force on the membranes to induce fusion as the complex ‘zippers’ from the N- to the C-terminus (10, 25, 29). The action of the SNAREs in this model can be rationalized in the framework of elastic continuum models of membrane fusion that are based on the effects of proteins on membrane properties such as curvature, elastic modulus and tension (30). These models postulate that membrane fusion occurs in several steps that are associated with large energy barriers and involve removal of hydration layers as the two membranes are brought together, bilayer bending to cause protrusions often called nipples, formation of a stalk intermediate in which the proximal leaflets have fused, merger of the distal leaflets to form a fusion pore, and opening of the fusion pore (30, 31). The textbook model was also supported by coarse-grained (CG) molecular dynamics (MD) simulations, which suggested that helix continuity is crucial for the SNAREs to cause fusion and that the mechanical forces arising from SNARE zippering not only bring the membranes together but also facilitate the downstream steps that lead to fusion (32–34).

**Figure 1.**
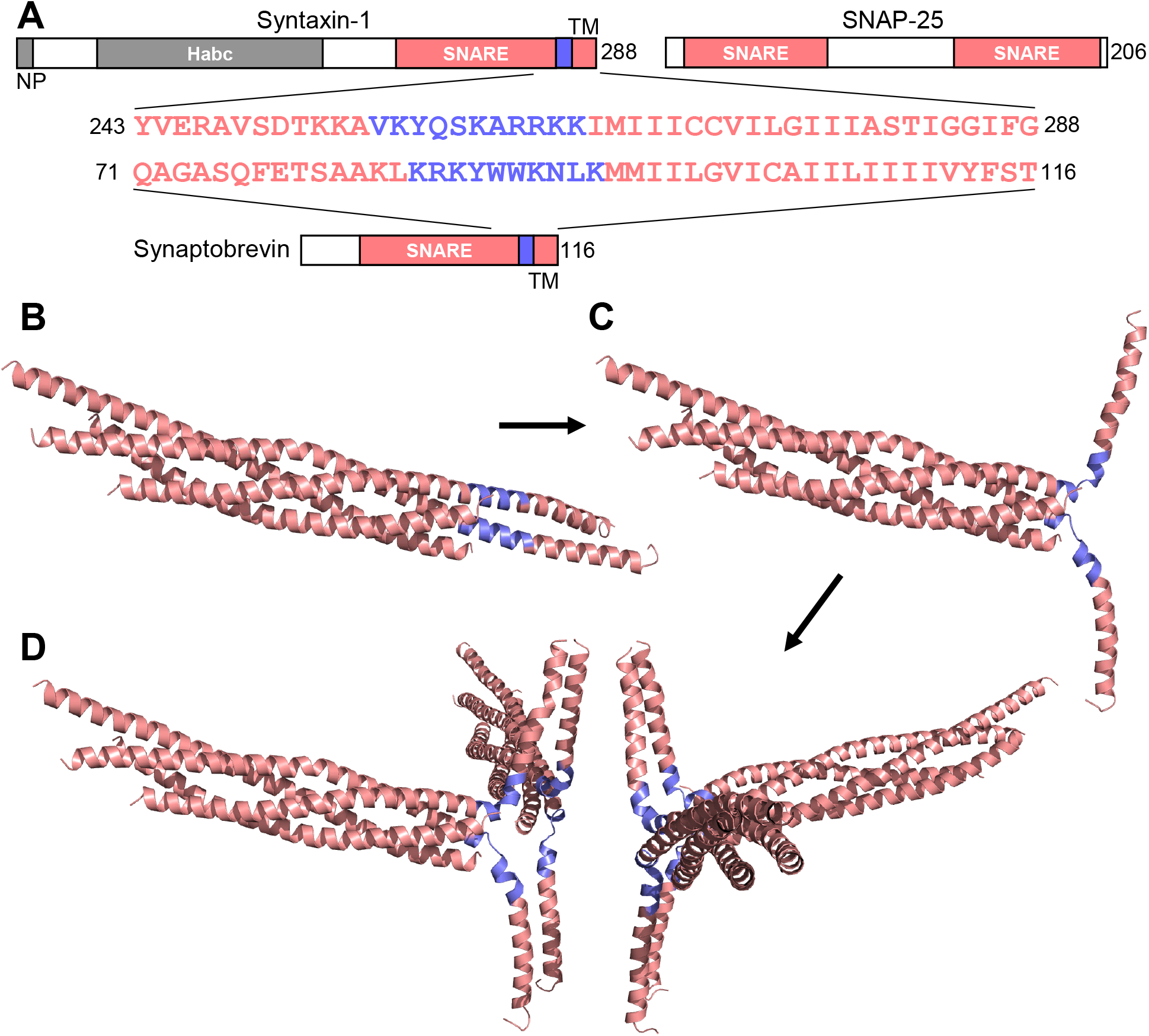
Generation of almost completely helical trans-SNARE complexes. (*A*) Domain sequences of syntaxin-1, SNAP-25 and synaptobrevin, with the SNARE motifs and TM regions in salmon, and the jxt linkers in blue. Numbers on the right of the diagrams show the length of each protein. NP = N-peptide; SNARE = SNARE motif; TM = transmembrane. The sequences of the C-terminal residues of the SNARE motifs, the jxt linkers and the TM regions of syntaxin-1 and synaptobrevin are shown. (*B*) Ribbon diagram of the crystal structure of the neuronal cis-SNARE complex (28) (PDB accession number 3HD7), which was used as starting point for restrained MD simulations used to generate trans-SNARE complexes. The N-terminal region of syntaxin-1 containing the N-peptide and the H_abc_ domain is not part of the structure and was not included in any of the MD simulations described here. (*C*) Ribbon diagram of the trans-SNARE complex generated by restrained MD simulations with mild force constants to minimal perturb the conformations of the jxt linkers. (*D*) Ribbon diagram of the four trans-SNARE complexes obtained by rotations and translations of the complex shown in (*C*), after they were incorporated in the fusion2g system and the temperature and pressure of the system were equilibrated. The jxt linkers are colored in blue in (*B-D*).

A considerably different picture emerged from CG MD simulations with two protein-free small (14-15 nm) vesicles placed in contact, which suggested that the key step to initiate membrane fusion is the encounter of acyl chains from lipids of the apposed bilayers at the polar interface, leading to formation of a small hydrophobic core that quickly progresses to form a stalk intermediate and later a fusion pore without the help of proteins (35–37). This overall model was supported by all-atom MD simulations of 15 nm protein-free vesicles (38), but no encounters between lipid acyl chains at the polar interface were observed in MD simulations with 14 nm vesicles using an improved CG force field (39). Interestingly, such encounters were facilitated by lung surfactant protein B, which bridged the two vesicles by binding to them through amphipathic α-helices, and fusion ensued quickly (< 500 ns) after the initial acyl chain encounter, with the protein bound only to the vesicle surface (39).

It is also important to note that insertion of short helix-breaking sequences between the synaptobrevin jxt and TM sequences still allows robust liposome fusion in vitro and granule exocytosis in chromaffin cells (40–42), and a similar insertion between the syntaxin-1 jxt and TM sequences still allows neurotransmitter release in neurons (43). These results showed that continuous helices in synaptobrevin and syntaxin-1 are not required for membrane fusion and neurotransmitter release. In contrast, insertion of short helix-breaking sequences between the SNARE motif and jxt linker of synaptobrevin or syntaxin-1 strongly impairs liposome fusion (42) or neurotransmitter release (43, 44), respectively, showing that zippering of the jxt linker does play a key role in SNARE action. However, the molecular basis for these observations and the overall mechanism of SNARE-induced membrane fusion remain unclear.

We have addressed these questions using all-atom MD simulations, taking advantage of the power of current supercomputers that allow the calculation of trajectories of a few microseconds for systems of millions of atoms, and building on our recent all-atom MD simulations of the neurotransmitter release machinery bridging a 24 nm vesicle and a flat lipid bilayer (23). Importantly, we observed fusion of the vesicle and the flat bilayer in less than 2 μs at 350 K in a simulation in which the jxt linkers of synaptobrevin or syntaxin-1 were fully zippered, whereas no fusion was observed without such zippering. While further simulations will be required to draw more firm conclusions, our results uncover a compelling molecular mechanism that makes a lot of sense from a physicochemical point of view and explains a large amount of experimental data. In this mechanism, membrane fusion depends more on local re-arrangements than on overall membrane properties, as membrane curvature did not appear to play an important role and no overt nipple formation was observed. Instead, the crucial event that initiated fusion was the catalysis of lipid acyl chain fluctuations out of the bilayers by the zippered linkers and adjacent hydrophobic residues from the TM regions. Encounters of the acyl chains from the two bilayers formed a hydrophobic core that quickly expanded into stalk-like structures and gradually progressed into a fusion pore, with the help of the TM regions and without going through a clearly defined intermediate. Based on other available data and the importance of the jxt linkers for other types of intracellular membrane fusion [e.g. (45, 46)] (see discussion), it seems likely that central aspects of this mechanism are generally conserved.

## Results

### Four trans-SNARE complexes with fully assembled four-helix bundles bridging a vesicle and a flat bilayer

The extremely fast speed of neurotransmitter release in fast synapses (27) suggests that synaptic vesicle fusion can be triggered in a few microseconds when the release machinery reaches proper configurations and hence that it may be possible to reproduce this process through all-atom MD simulations. However, the minimum number of trans-SNARE complexes required for fusion and their locations in these configurations are uncertain. In a previous study (23) we performed simulations of a 24 nm vesicle and a flat lipid bilayer bridged by four trans-SNARE complexes that were placed at the periphery of the bilayer-bilayer interface and were generated by an initial restrained MD simulation started with the crystal structure of the cis-SNARE complex (28). During the initial restrained simulation, the TM regions of syntaxin-1 and synaptobrevin quickly moved to designed target positions and remained helical, and the SNARE four-helix bundle remained almost fully assembled, but the jxt linkers became unstructured. This was not surprising, as continuous helical conformations throughout the entire syntaxin-1 and synaptobrevin sequences would have required unrealistic bending of the helices and dissociation of part of the four-helix bundle, which is very stable (47). During the simulation of the whole system, the vesicle and the flat bilayer quickly formed an extended interface that resembled those observed in cryo-EM images of SNARE-mediated liposome fusion reactions (26), and no fusion was observed after 520 ns at 310 K and 454 ns at 325 K (23).

To explore potential SNARE configurations that might induce fast membrane fusion, we reasoned that placing the trans-SNARE complexes closer to each other may facilitate the formation of nipples, which is postulated to occur before fusion (30). Moreover, proximity to the center of the bilayer-bilayer interfaces is expected to facilitate the retention of helical conformation in the jxt linkers, which may also facilitate fusion. To generate trans-SNARE complexes for this purpose, we also started with the crystal structure of the cis-SNARE complex (28) and performed a 2 ns restrained MD simulation in which the TM regions were targeted to designed positions using a mild force constant (10 kJ*mol^−1^*nm^−2^) to try to perturb the helical structure as little as possible. In addition, we carried out a 1 ns restrained simulation with a slightly stronger force constant (50 kJ*mol^−1^*nm^−2^) to force the TM regions closer to the desired positions. The resulting trans-SNARE complex was almost completely helical except for kinks in the helices at the jxt linkers (Fig. 1*C*). We made four copies of this complex and placed them in designed positions with rotations and translations (Fig. 1*D*). The full system was built with these four complexes, a flat lipid bilayer (a square of 30.5 nm x 30.5 nm) built at the CHARMM-GUI website (48) (https://charmm-gui.org/) and the same 24 nm vesicle that we used previously (23) with several lipids moved manually to accommodate the different positions of the synaptobrevin TM regions. The lipids compositions of the vesicle and flat bilayer resembled those of synaptic vesicles and synaptic plasma membranes, respectively (49, 50), and the system (referred to as fusion2g) had 4.4 million atoms after solvation with water (Table 1).

**Table 1.**
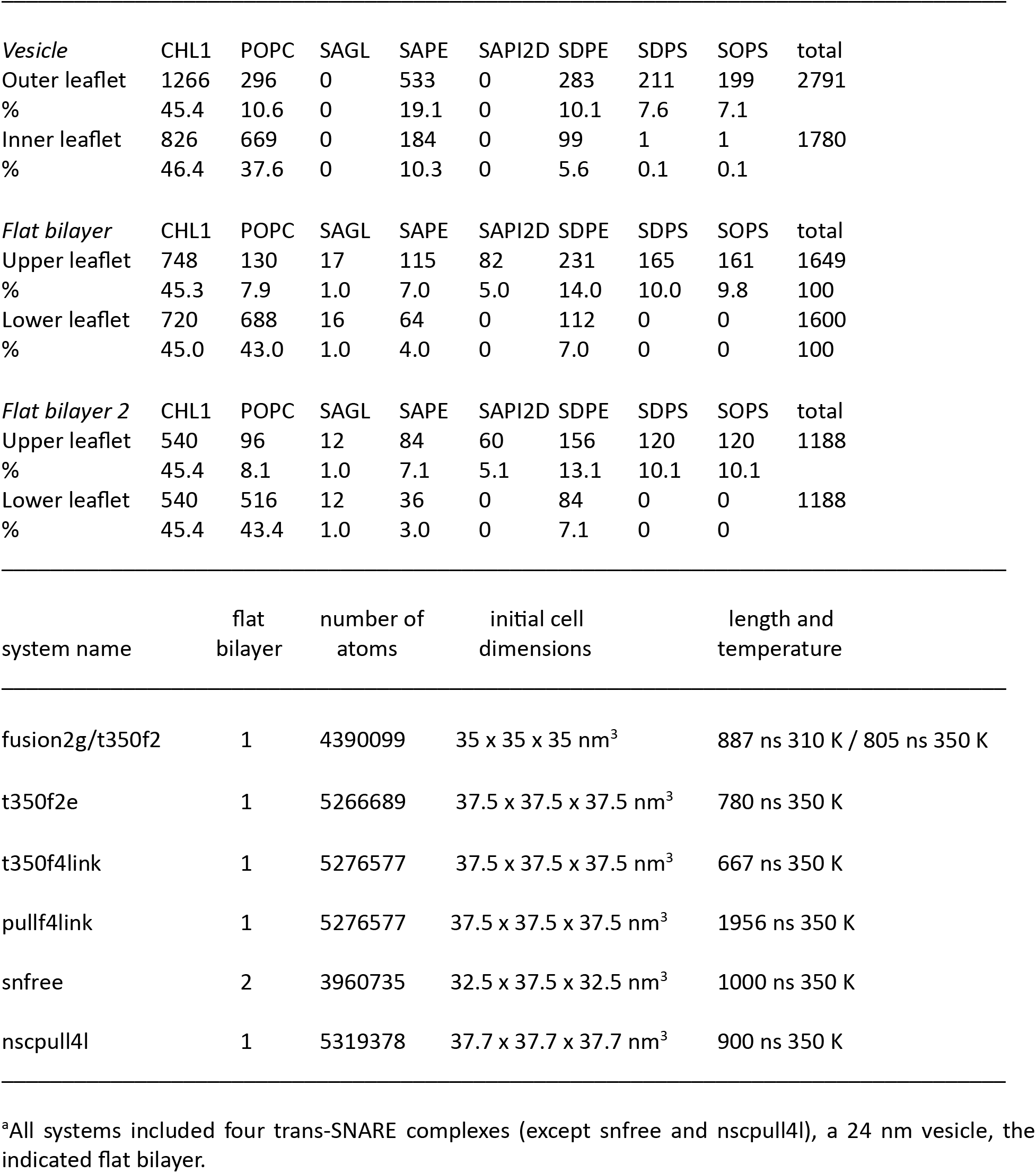
Parameters of the MD simulations^a^.

After equilibrating this system, we run a production simulation at 310 K for 857 ns. During the first 180 ns, the flat bilayer buckled slightly at the center, between the SNARE helices, but the buckling did not increase and there was no initiation of fusion at the end of the simulation (Fig. 2*A-C*). Since elevated temperatures can help to overcome energy barriers and are often used in MD simulations of lipid bilayers to increase lipid fluidity [e.g. (32)], we used the final configuration of the fusion2g simulation as a starting point to run another MD simulation at 350 K (referred to as t350f2). Due the natural rotation of the system that had already been occurring during fusion2g and continued in t350f2, part of the edge of the flat bilayer came out at the bottom of the cell used for periodic boundary conditions and emerged at the top of the cell, making contact with the vesicle early in the simulation (around 60 ns; Fig. 2*D*). As the simulation progressed, the edge of the bilayer merged with the top of the vesicle (Fig. 2*E*) and the flat bilayer gradually adopted a semispherical shape to adapt to the vesicle shape as the merger of the two bilayers extended (Fig. 2*F*). The merger of the two bilayers was initiated by one 1-stearoyl-2-docosahexaenoyl-sn-glycero-3-phosphoethanolamine (SDPE) molecule from the flat bilayer, which splayed when the polyunsaturated acyl chain moved into the vesicle outer leaflet (Fig. S1*A*). This SDPE molecule remained splayed for more than 200 ns and initiated bilayer merger only when its motions facilitated the movement of other acyl chains to the polar interface between the two bilayers, forming a small hydrophobic nucleus that quickly expanded in less than 20 ns (Fig. S1*B-H*).

**Figure 2.**
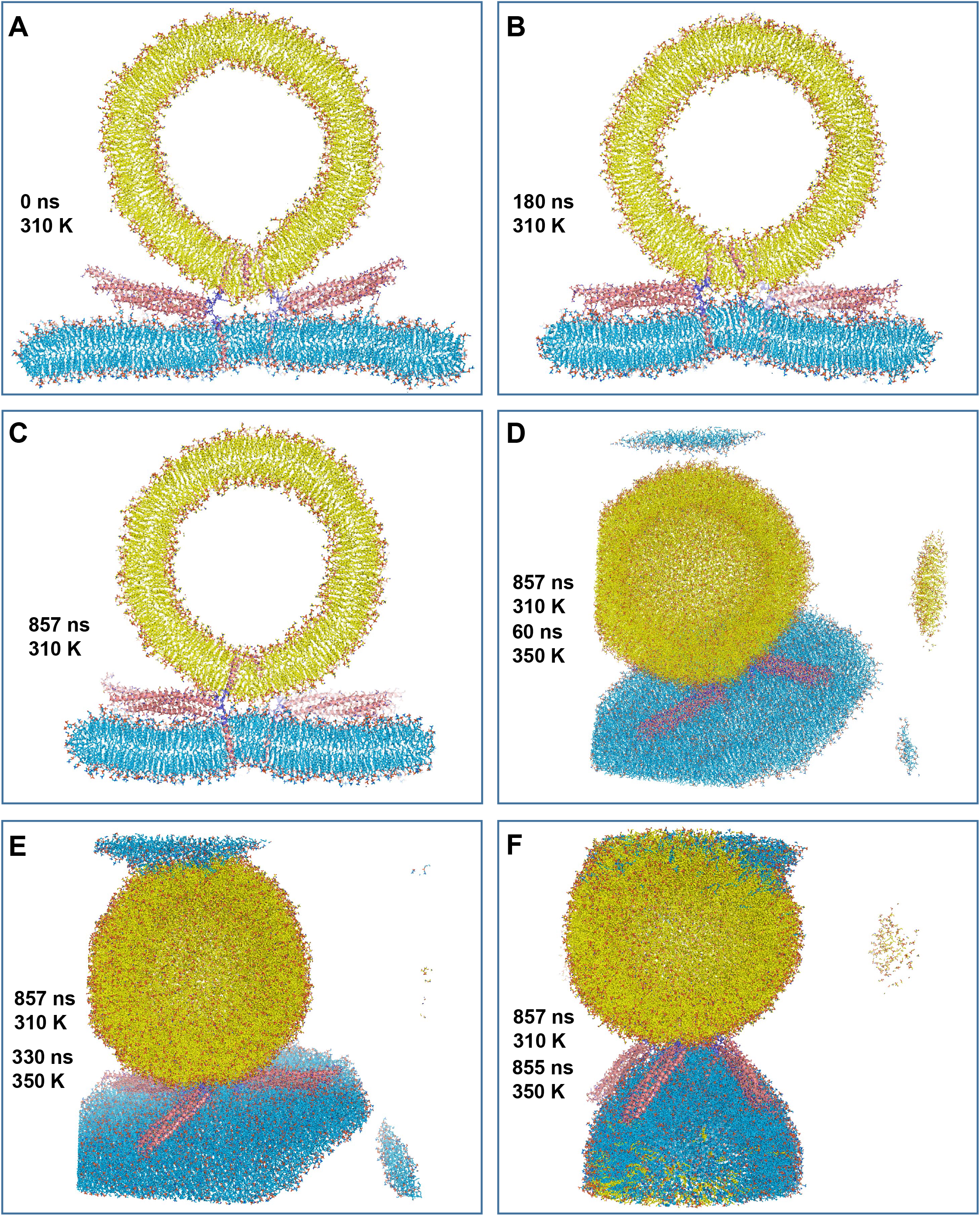
MD simulation of four almost completely helical trans-SNARE complexes bridging a vesicle and a flat bilayer and close to the center of the interface. (*A-C*) Slices of the system in its initial configuration (*A*) and after 180 ns (*B*) or 857 ns (*C*) of simulation at 310 K. The systems of (*B*) and (*C*) were superimposed with the initial configuration to facilitate comparisons. (*D-F*) Overall view of the system after 857 ns of simulation at 310 K and 60 ns (*D*), 330 ns (*E*) and 855 ns (*F*) at 350 K. The systems were not superimposed with the initial configuration so that the effects of random translation and rotation can be observed. The parts of the system that moved out of the box used for periodic boundary conditions emerge at the opposite side of the box. The emergence of the edge of the flat bilayer at the top of the box led firs to contact with the vesicle (*D*) and later to merger of the two bilayers (*E, F*). In all panels, lipids and proteins are shown as stick models with nitrogen atoms in dark blue, oxygens in red, phosphorus in orange and carbon atoms in yellow for the vesicle, light blue for the flat bilayer and salmon for the SNARE complex except for those of the jxt linkers, which are in purple. The same color-coding was used in all the figures except when noted otherwise. The SNARE complexes are in addition represented by ribbon diagrams.

Although the observed membrane merger was an artifact arising from the rotation and the insufficient thickness of the water layer surrounding the system, it is interesting that the mechanism of merger is reminiscent of that observed previously in CG and all-atom simulations with 14-15 nm protein free vesicles (35–38). In contrast, we did not observe even the initiation of membrane fusion by the SNAREs at the opposite side of the vesicle during the 857 ns at 310 K and the 855 ns at 350 K of the two simulations. To test the possibility that the lack of SNARE-mediated membrane fusion might have arisen because of the changes in shape of the flat bilayer as it was adapting to the vesicle shape, we took the coordinates of the system at 60 ns of the t350f2 simulation, before the membranes merged, re-oriented and centered the system in the cell, and built a larger water box to prevent contact between the edge of the flat bilayer and the vesicle due to system rotation. We ran a 780 ns MD simulation at 350 K of this system (referred to as t350f2e) but again did not observe even the initiation of SNARE-induced membrane fusion (Fig. S2). Indeed, we did not observe the initiation of membrane fusion between a flat bilayer and a 24 nm vesicle brought into direct contact by fully assembled or close to fully assembled SNARE four-helix bundles in multiple MD simulations that we reported previously (23), that are discussed here or will be described elsewhere, and that altogether totaled over 15 μs. Although, admittedly, this is still a short time scale, these findings correlate with the observation of that extended contact interfaces between liposomes induced by neuronal SNARE complexes and observed by cryo-EM fuse slowly, in the second-minute time scales (26), and suggest that the packing of the lipids in the flat bilayers and the 24 nm vesicles of our simulations is tight enough to strongly hinder the movement of the acyl chains to the polar interface to form non-bilayer structures. Conversely, the observation of such movement and subsequent membrane merger in our t350f2 simulation most likely arises because of poor lipid packing at the edge of the flat bilayer, which facilitated the splaying of the SAPE molecule (Fig. S1). Similarly, the fusion between 14-15 nm protein-free vesicles observed previously (35–38) may have arisen because of their strong curvature, which also favors poor lipid packing.

### Jxt linker zippering leads to fast vesicle-flat bilayer fusion

The functional consequences of inserting short helix-disrupting sequences before or after the jxt linker of synaptobrevin or syntaxin-1 showed that a continuous helix including the SNARE motif, jxt linker and TM sequence is not required for liposome fusion in vitro or neurotransmitter release in neurons, but extension of the SNARE motif helices into the jxt linkers (referred to as jxt linker zippering) is actually crucial (41–44). Our MD simulations suggest that a substantial energy barrier hinders such extension because of the natural geometry of trans-SNARE complexes, which dictates that the angle between the SNARE four-helix bundle and the TM regions is much smaller than 180° (close to 90° for the geometry of the fusion2g system; Fig. 1), and leads to unstructured (23) or kinked-helical conformations for the jxt linkers. However, zippering of the jxt linker is associated with a substantial folding energy [8 k_B_T (47)] that can help overcome this energy barrier and may occur spontaneously, albeit slowly, and/or may be favored by other components of the neurotransmitter release machinery.

Since jxt linker zippering may be a slow step that cannot be reached naturally in the time scales that we can simulate, we used restrained MD simulations to induce such zippering and investigate its potential effects on membrane fusion. Since the TM regions of the trans-SNARE complexes of the fusion2g simulation were quite close to each other and jxt linker zippering would bring them even closer, we first moved the trans-SNARE complexes manually from the fusion2g simulation outward, i.e. further from each other, and used restrained MD simulations to gradually move the TM regions to new designed positions. This procedure led to stretching of the conformations of the jxt linkers because the TM regions of synaptobrevin and syntaxin-1 are located further from each other in the new locations (Fig. 3*A*). We then used additional restrained MD simulations to induce helical conformations in the jxt linkers, up to residue 94 of synaptobrevin and 265 of syntaxin-1, using the structure of the cis-SNARE complex (28) as a model (Fig. S3*A*). The trans-SNARE complexes were merged with the same flat bilayer and vesicle used for the fusion2g simulation, and lipids of both membranes were moved manually to accommodate the new TM positions. After equilibrating the system first at 310 K and then at 350 K, we ran a production simulation at 350 K for 667 ns (referred to as t350f4link; Fig. S3*B-F*). The two membranes were brought into close contact during the simulation and some of the TM regions came close to each other, buckling the flat bilayer a little bit (Fig. 3*C-F*), but there was no initiation of fusion. The jxt linkers remained partially helical throughout the simulation, with some oscillations in the degree of helicity, and the helices were separated at the C-termini to different extents by the end of the simulation (Fig. S3*B*).

**Figure 3.**
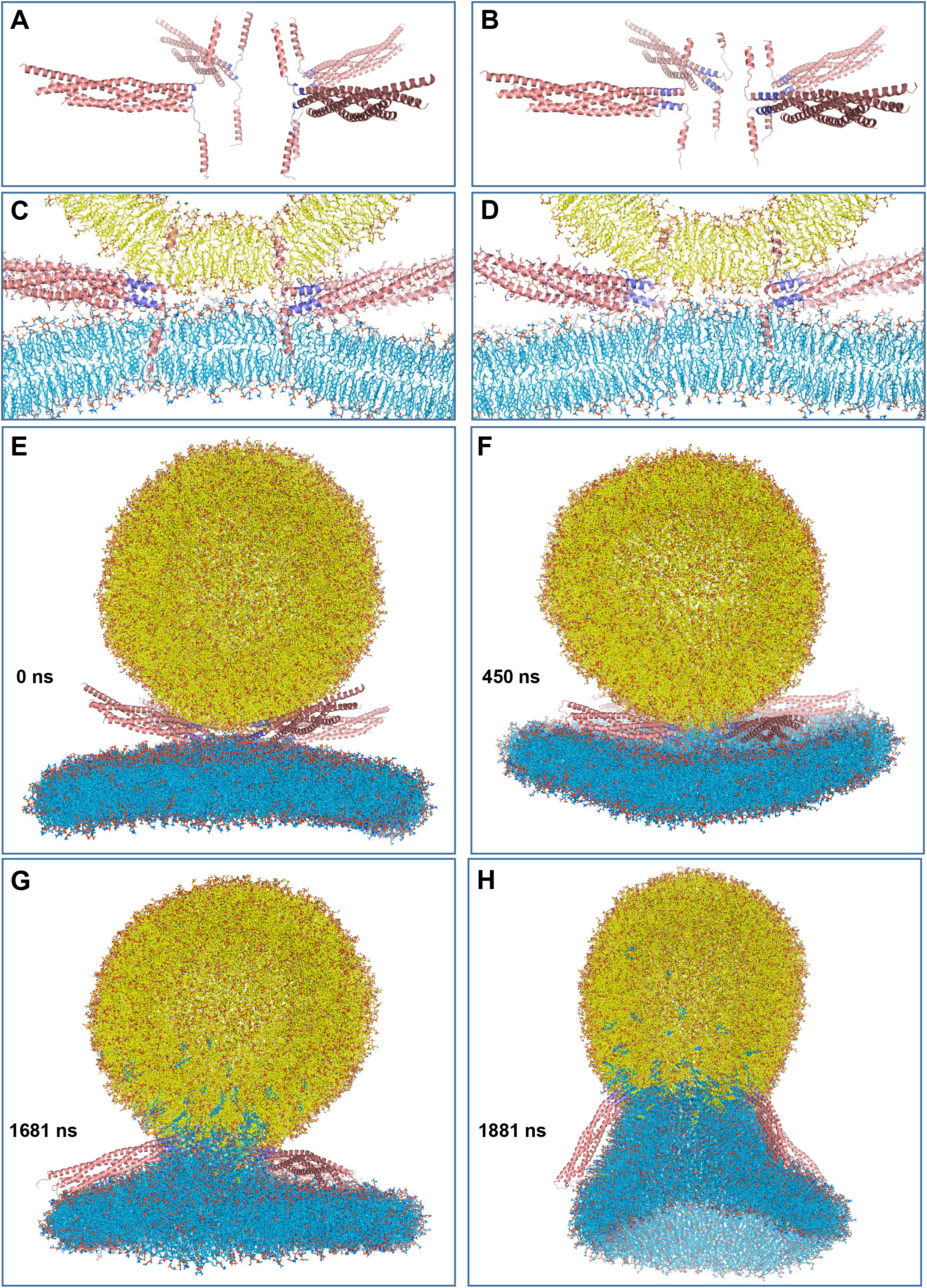
MD simulation of four trans-SNARE complexes bridging a vesicle and a flat bilayer, with the linkers zippered and a pulling force to keep them zippered. (*A-B*) Ribbon diagrams showing initial configuration of the trans-SNARE complexes before using restrained MD simulations to zipper the linkers (*A*) and at the end of these simulations, with the linkers fully zippered (*B*). (*C-D*) Slices of the system showing the trans-SNARE complexes inserted into the flat bilayer and the vesicle in the initial configuration used for the pullf4link simulation, showing from opposite angles to display the four complexes. (*E-H*) Full views of the system at the indicate time points of the pullf4link simulation. Lipids and proteins are shown as stick models, and SNARE complexes are in addition represented by ribbon diagrams. The color-code is the same as in Fig. 2.

To facilitate permanent full zippering of the linkers, we generated a similar system in which the trans-SNARE complexes were induced to adopt helical conformation a little beyond the linkers (up to residue 99 of synaptobrevin and 268 of syntaxin-1) using also the cis-SNARE complex structure (28) as a model (Fig. 3*B*) and the initial equilibrated configuration of the t350f4link system as the starting point (i.e. including the flat bilayer and the vesicle). Compared to the trans-SNARE complexes before forcing any helical conformations in the linkers (Fig. 3*A*), this procedure brought the TM regions of synaptobrevin molecules closer to the syntaxin-1 TM regions, as expected, and stretched them such that their N-termini became partially unstructured (Fig. 3*B*). Correspondingly, the C-termini of synaptobrevin and syntaxin-1 were brought inside their corresponding bilayers (Fig. 3*C-D*). We then ran a 100 ns production simulation at 310 K followed by 1956 ns at 350 K while using the pull code from Gromacs (51, 52) to restrain the distance between the geometric centers of the atoms of residues 94-97 of synaptobrevin and of the atoms of residues 263-266 of syntaxin-1 to within 1 nm, which is their distance in the cis-SNARE complex structure (referred to as pullf4link simulation).

Importantly, as illustrated by full views of the system in Fig. 3*E-H* and the system slices of Fig. 4, this simulation did led to full fusion of the vesicle and the flat bilayer. The two membranes were in close contact early in the simulation and retained their bilayer structures until about to 400 ns (Fig. 4*B*). An stalk-like structure began to form shortly afterwards (Fig. 3*F*, 4*C*) and lipids at this region kept re-arranging (e.g. Fig. 4*D*) during over 1000 ns, leading to full merger of the two bilayers to initiate formation of a fusion pore (Fig. 3*G*, 4*E*) that opened very quickly (Fig. 3*H*, 4*F*).

**Figure 4.**
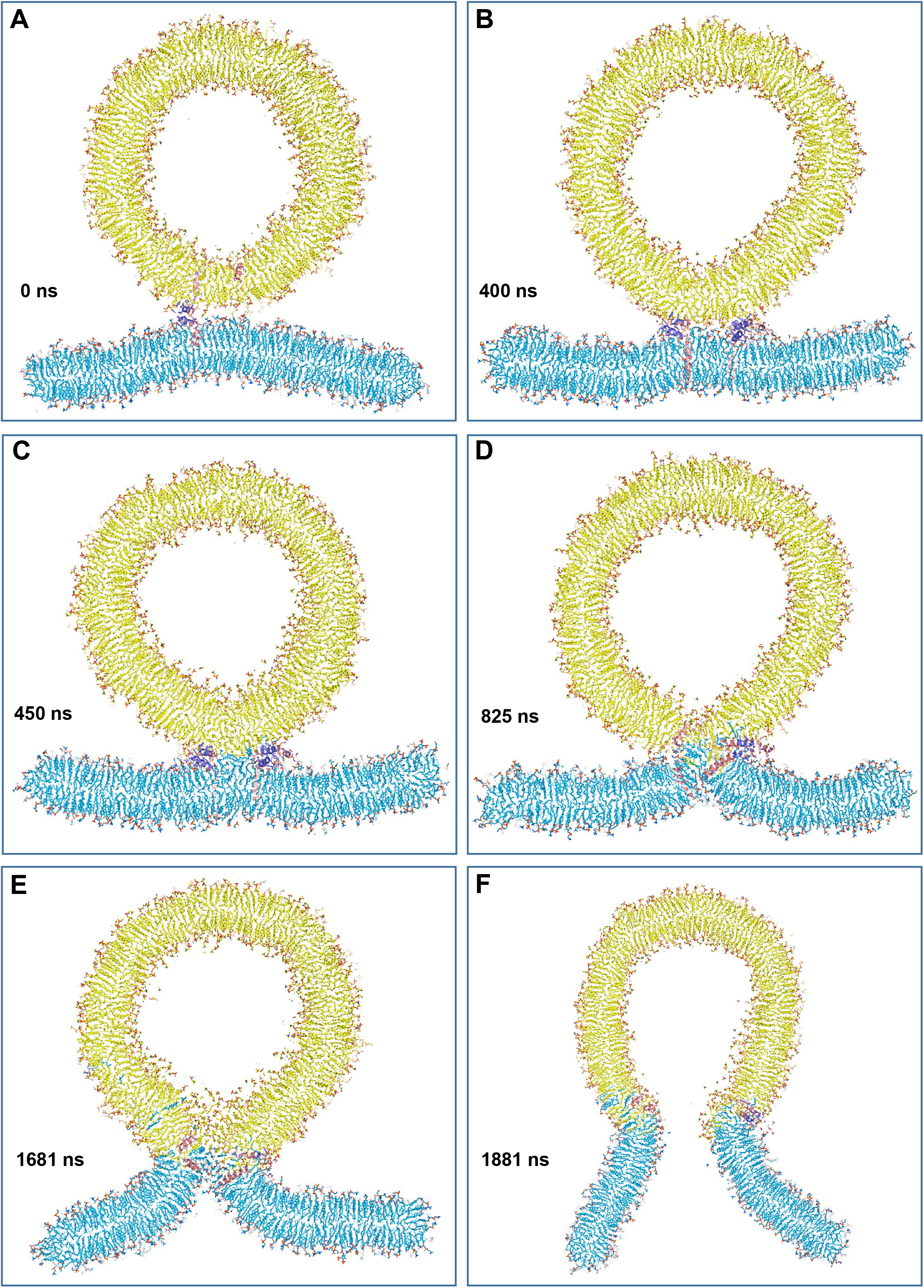
Slices taken at the indicated time points along the pullf4link simulation. The slices illustrate how the membranes retained their bilayer structure from 0 to 400 ns (*A,B*), a small stalk-like structure had formed by 450 ns (*C*), and the interface kept evolving (*D*) until a fusion pore was formed (*E*) and expanded (*F*). Lipids and proteins are shown as stick models, and SNARE complexes are in addition represented by ribbon diagrams. The color-code is the same as in Fig. 2.

### Mechanism of vesicle-flat bilayer fusion

Zippering of the linkers was crucial to initiate the merger of the vesicle and the flat bilayer because it brought the hydrophobic residues of the TM regions to the polar interface, where they facilitate the transition of hydrophobic acyl chains out of the two bilayers. In the trajectory it became apparent that these transitions are particularly likely for lipids containing polyunsaturated acyl chains that are abundant in the vesicle and plasma membranes (49, 50). The vesicle and flat bilayer contained three types of such lipids (number of carbons and unsaturated bonds indicated in parenthesis): SDPE (18:0, 22:6), 1-stearoyl-2-arachidonoyl-sn-glycero-3-phosphoethanolamine (SAPE; 18:0) and 1-stearoyl-2-docosahexaenoyl-sn-glycero-3-phospho-L-serine (SDPS; 18:0, 22:6). Figure 5 illustrates how the 22:6 acyl chain of an SDPS molecule started to emerge from the flat bilayer already at 75 ns and was fully splayed by 80 ns. These events were clearly facilitated by contacts of this acyl chain with the hydrophobic side chains of the TM regions and the methylene groups of lysine side chains from the jxt linkers of one SNARE complex. However, there was no initiation of lipid mixing in this area of the vesicle-flat bilayer interface, suggesting that lipid splaying itself is not sufficient to induce membrane merger.

**Figure 5.**
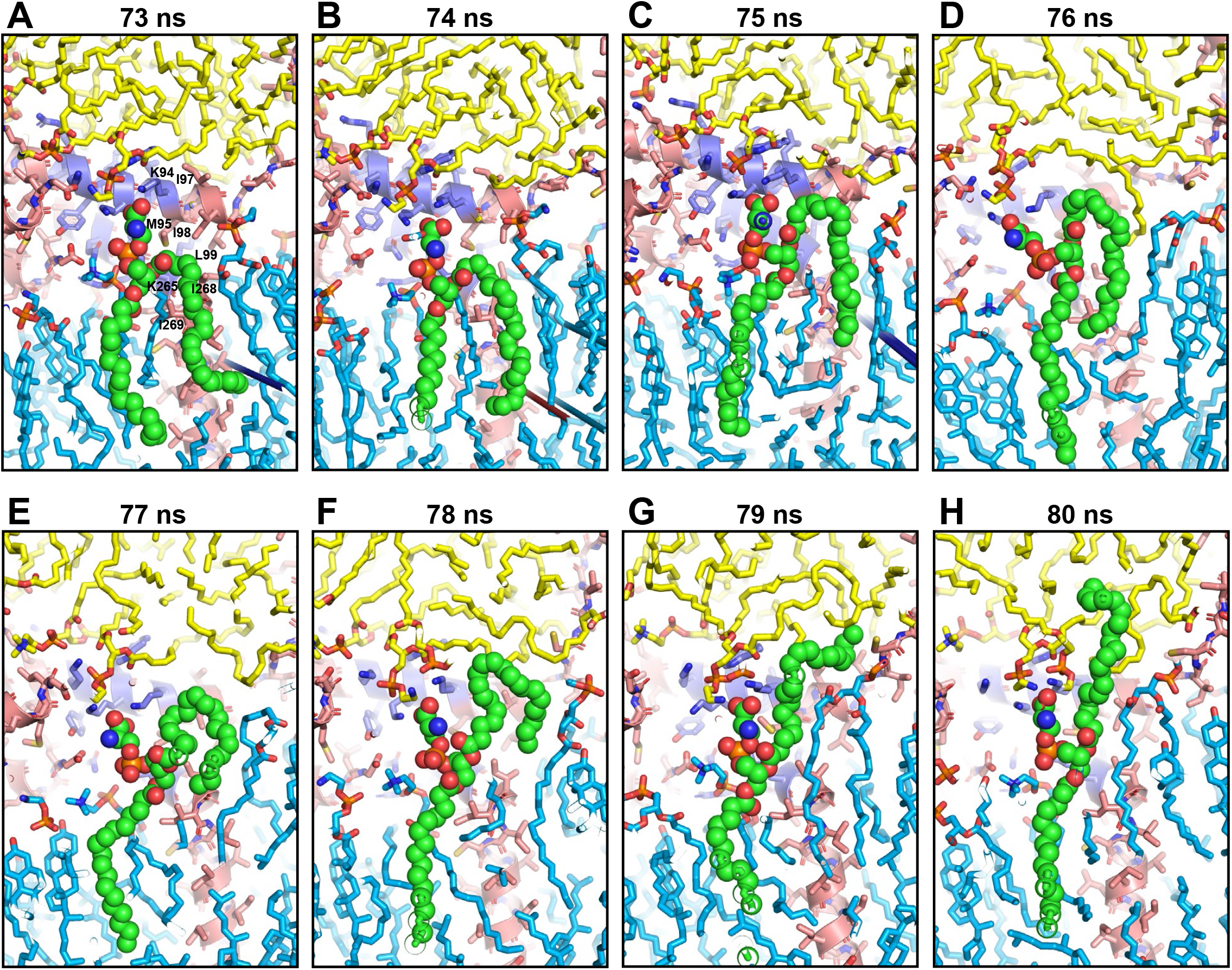
Snapshots taken at the indicated time points of the pullf4link simulation showing the splaying of an SDPS molecule catalyzed by the jxt linkers and TM regions of one of the trans-SNARE complexes. Lipids and proteins are shown as stick models, and SNARE complexes are in addition represented by ribbon diagrams. The color-code is the same as in Fig. 2. The SDPS molecule is represented by spheres with nitrogen atoms in dark blue, oxygens in red, phosphorus in orange and carbon in green.

Membrane merger was initiated by another SNARE complex (Fig. 6) when an acyl chain from an SDPS molecule from the flat bilayer (2501) emerged at the polar interface and made contact with acyl chains of two SAPE molecules from the vesicle (8037 and 8045) that also emerged at the interface at 402 ns (Fig. 6*B*). All these acyl chains made contacts with hydrophobic side chains of the TM regions and methylene groups of lysine side chains from the jxt linkers of this SNARE complex. The acyl chain of an SDPE molecule from the flat bilayer (2012) soon emerged to join these hydrophobic contacts with the vesicle (Fig. 6*C*) and additional hydrophobic contacts were formed quickly at the interface (Fig. 6*D*) to initiate the formation of a small stalk-like structure in which the proximal leaflets of the flat bilayer and the vesicle merged. This mini stalk occurred in one side of the flat bilayer-vesicle interface (on the right in the orientation of Fig. 7*A*) and gradually expanded toward the left on this orientation (Fig. 7*B*), but polar groups from both bilayers remained at the interface and a full stalk spanning the entire interface never formed. Lipid re-arrangements continued and led to structures in which the vesicle formed a V-shape and the flat bilayer formed an inverted V-shape as expected for the stalk intermediate (30) (Fig. 7*C, D*). These shapes helped to minimize the volume of the hydrophobic area at the center of the interface and the difficulty of properly packing the lipids in these non-bilayer structures. The lipids at the center of these structures were somewhat disordered and some polar groups remained at the hydrophobic interface, likely because they can interact with the backbone amide groups of residues from the synaptobrevin TM regions that are not helical (Fig. 7*C, D*). The non-symmetrical nature of these structures is illustrated by the views from different angles of the pose obtained at 1125 ns (Fig. 7*D-F*).

**Figure 6.**
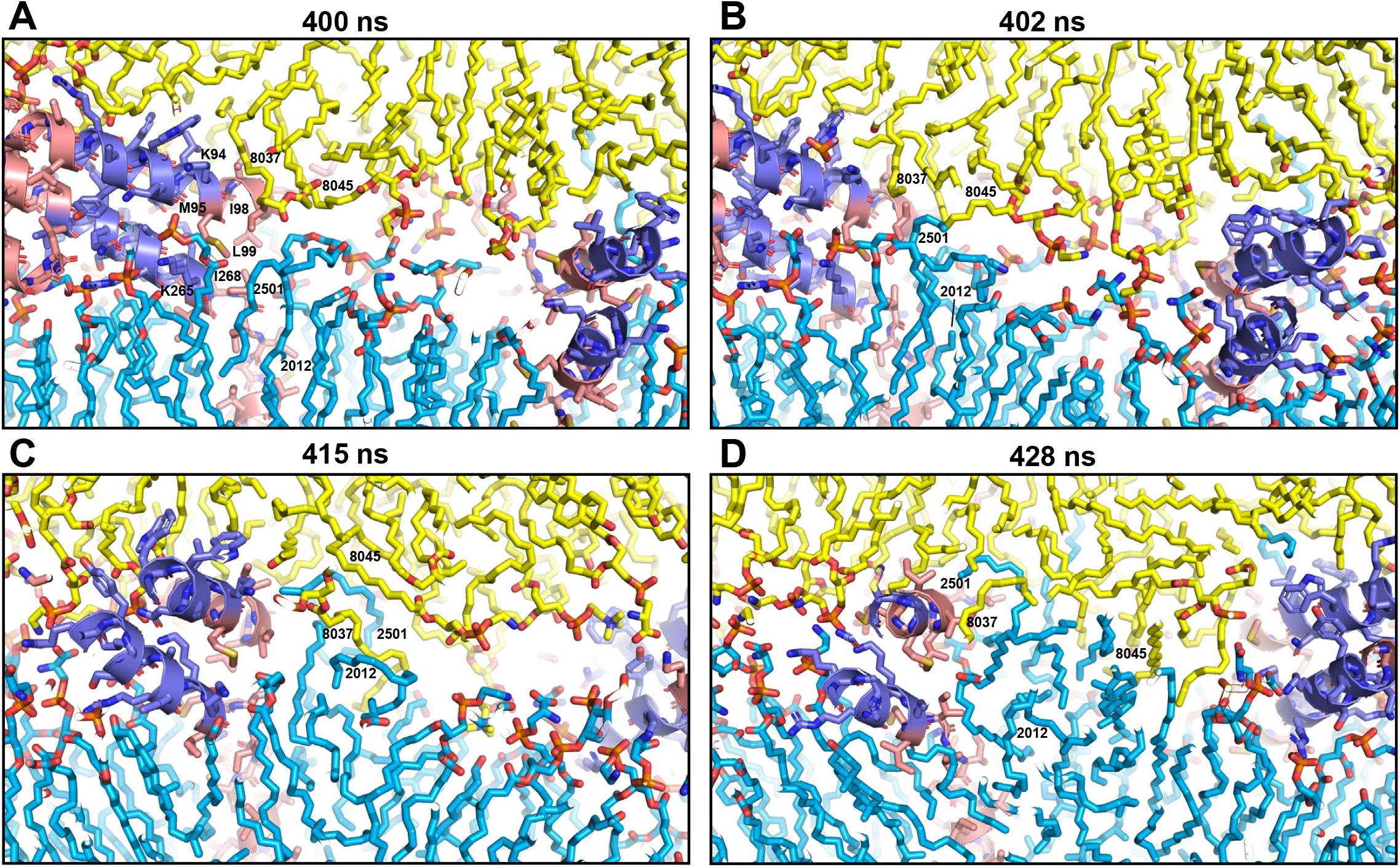
Snapshots taken at the indicated time points of the pullf4link simulation showing the formation of a small hydrophobic nucleus at the polar interface. The diagrams illustrate how there were no hydrophobic contacts in this region of the polar interface between the two bilayers at 400 ns (*A*). Acyl chains of polyunsaturated lipids from the two bilayers came into contact at the polar interface at 402 ns (*B*), when they came out of the bilayers due to fluctuations next to the methylene groups of lysine side chains from the linkers and hydrophobic side chains from the TM regions [labeled with the single letter amino acid residue abbreviation and residue number in (*A*)]. Additional contacts kept forming next to these residues (*C*) and facilitated the formation of hydrophobic contacts, forming a small hydrophobic nucleus (*D*). Lipids and proteins are shown as stick models, and SNARE complexes are in addition represented by ribbon diagrams. The color-code is the same as in Fig. 2. The positions of SDPE2012, SDPS2501, SAPE8037 and SAPE8045 are indicated to illustrate the movements of the lipids.

**Figure 7.**
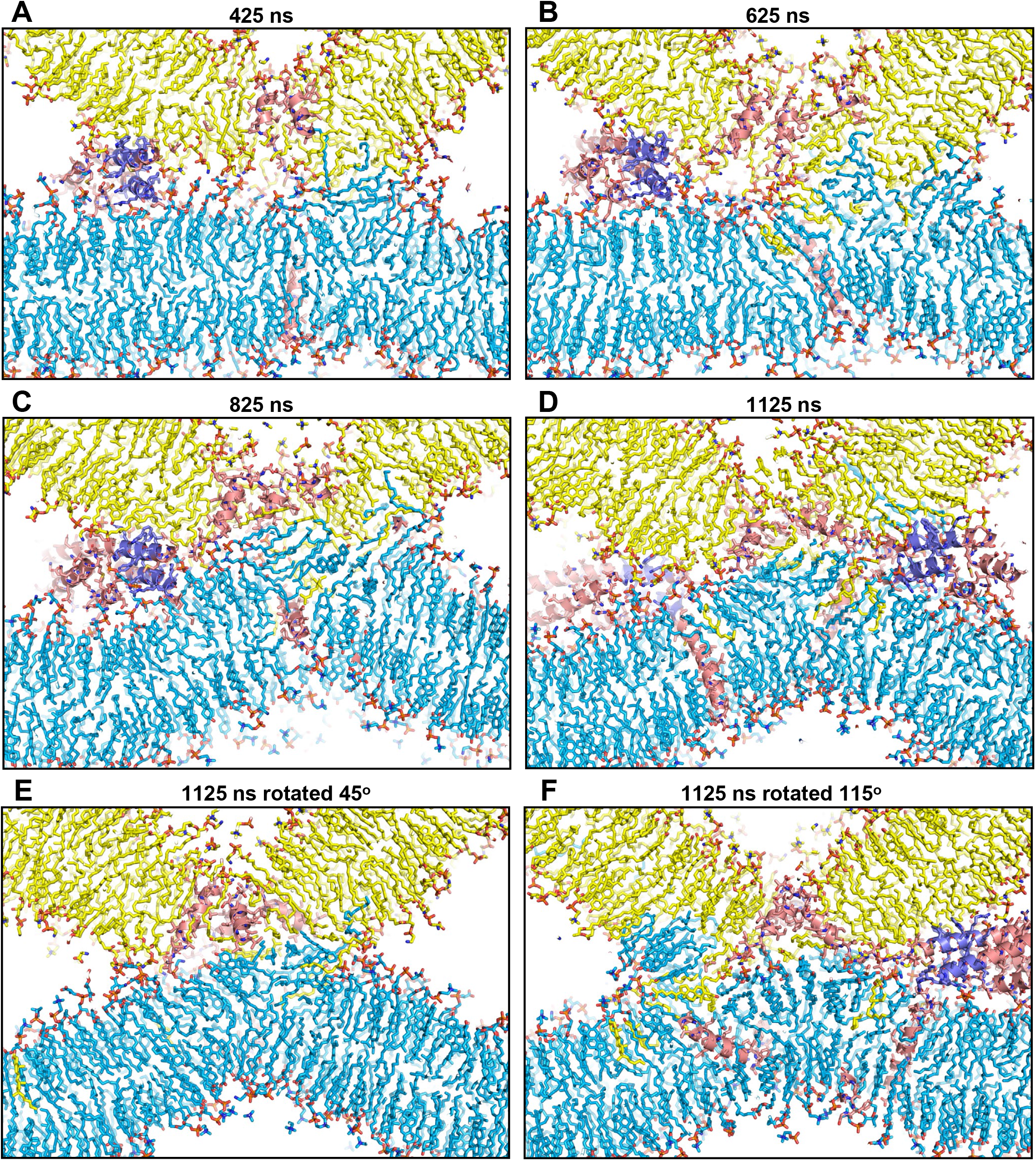
Snapshots of the pullf4link simulation showing the expansion of the hydrophobic nucleus into stalk-like structures. Panels (*A-D*) were taken at the indicated time points. Panels (*E,F*) show the same frame as (*D*) (at 1125 ns) but from different angles to illustrate the asymmetry of the lipid configuration at the interface, with some areas being purely hydrophobic while other areas contain lipid head groups that are normally close to the backbone of unstructured residues of the TM regions. Lipids and proteins are shown as stick models, and SNARE complexes are in addition represented by ribbon diagrams. The color-code is the same as in Fig. 2.

Polar lipid head groups gradually moved toward the center of the interface (Fig. 8*A-D*) until two bilayers that resulted from merging the flat bilayer and the vesicle were clearly formed, resulting in a nascent fusion pore (Fig. 8*E*) that quickly expanded (Fig. 8*F*, 4*F*). This quick expansion released the high curvature existing at the neck of the pore and may have been facilitated by the geometry of our system, as the small flat bilayer readily adapted to the shape of the vesicle (Fig. 3*H*), much as the flat bilayer accommodated to the shape of the vesicle when they merged in the t350f2 simulation (Fig. 2*F*). The movement of polar head groups to the center of the interface again appeared to be facilitated by interactions of the head groups with backbone amide bonds from unstructured residues of the synaptobrevin TM regions, which were interspersed with the lipids at the interface throughout the entire process of merging the membranes, as were the syntaxin-1 TM regions (Fig. 6-8). Interestingly, conformational flexibility in the synaptobrevin TM region has been shown to be crucial for efficient Ca^2+^- triggered exocytosis (53, 54). These observations correlate with the fact that, while the C-terminal half of the synaptobrevin TM regions (residues 106-116) were helical throughout the simulation, residues 100-105 remained largely unstructured (Fig. 9).

**Figure 8.**
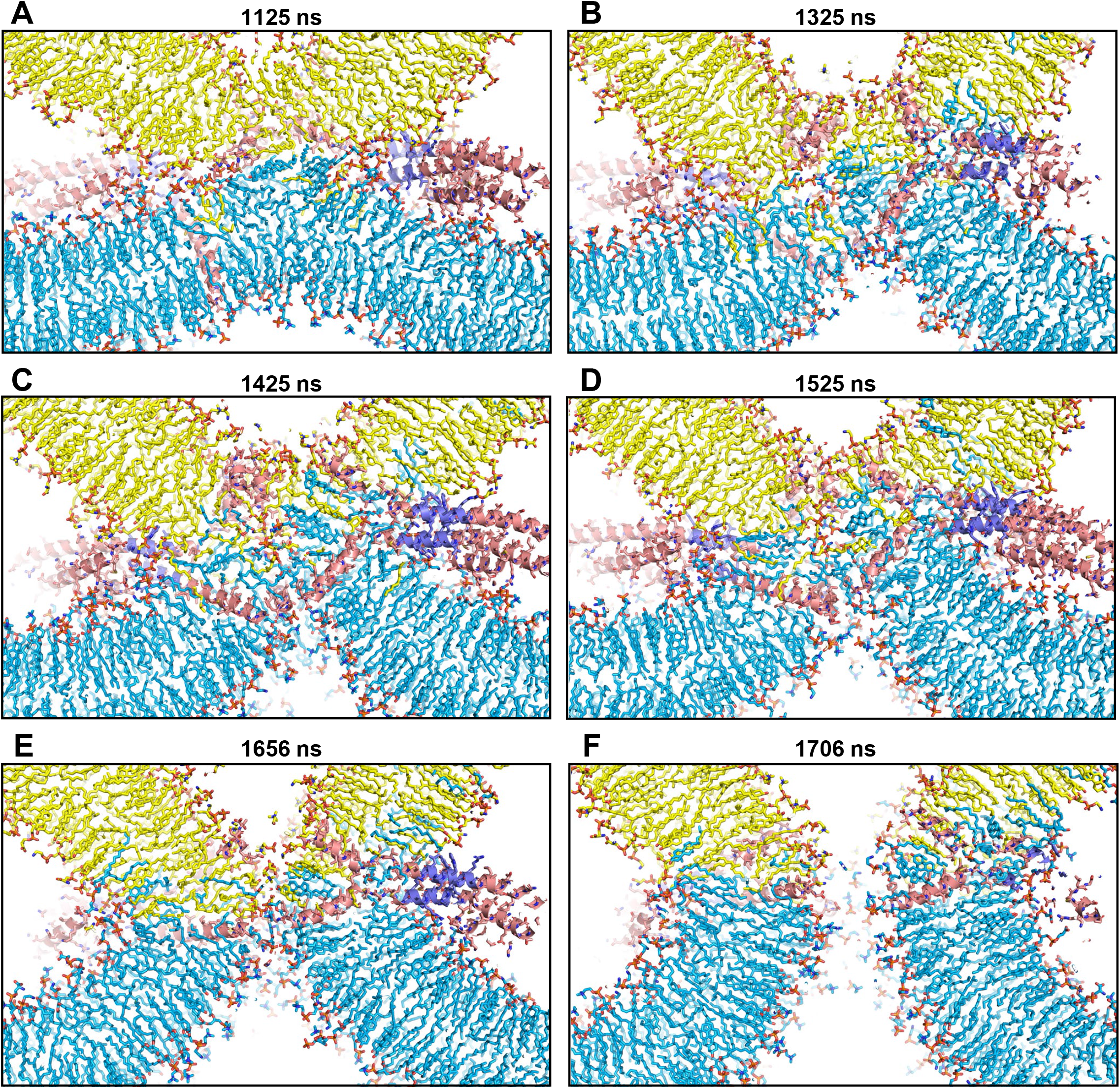
Snapshots taken at the indicated time points of the pullf4link simulation showing the evolution from stalk-like structures to the fusion pore. The diagrams show how the lipids were often quite disordered at the interface of the stalk-like structures and they became more ordered as they adopted the orientations that correspond to their positions in the developing fusion pore, most likely guided at least in part by the SNARE TM regions. Lipids and proteins are shown as stick models, and SNARE complexes are in addition represented by ribbon diagrams. The color-code is the same as in Fig. 2.

**Figure 9.**
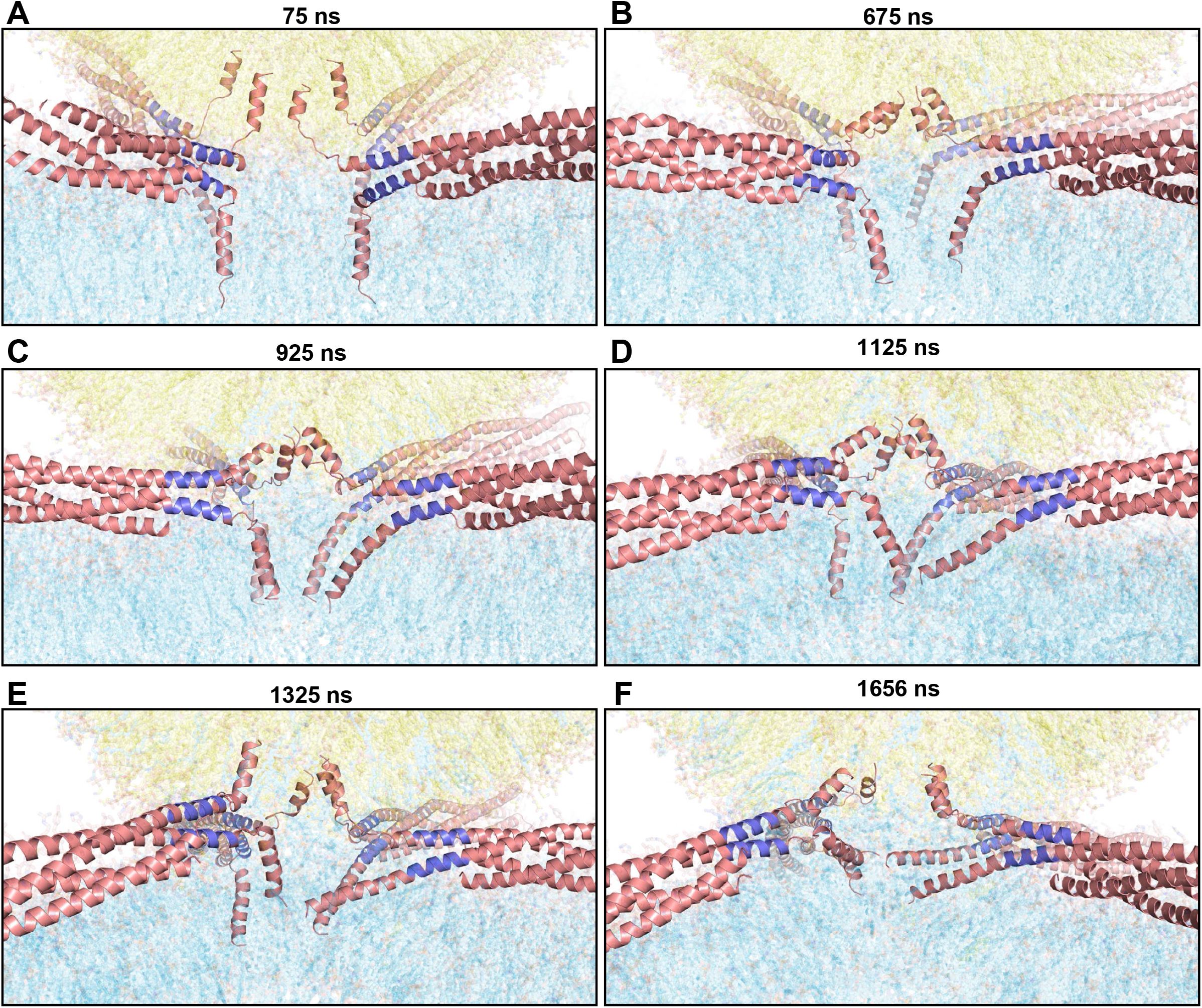
Snapshots taken at the indicated time points of the pullf4link simulation illustrating the conformational changes in the SNAREs as the system evolved from bilayer-bilayer contact to the fusion pore. SNARE complexes are represented by ribbon diagrams and lipids are represented by 90% transparent sticks to allow visualization of the sticks. Note that the thickness of the slices shown in these panels is much larger than those of the slices shown in Figures 4-8 so that all four trans-SNARE complexes can be observed, but the lipids then need to be transparent to allow visualization of the SNAREs. The color-code is the same as in Fig. 2.

A substantially different behavior was observed for the syntaxin-1 TMs. Although the N-terminal regions of the TMs, close to the jxt linkers, became unstructured after the restrained MD simulation used to set up the initial configuration of this system (Fig. 3*B*), they quickly regained most of the helicity (Fig. 9*A*). Two of the syntaxin-1 TM regions actually became fully helical by 675 ns, with a slight kink so that the helices were mostly parallel to the flat bilayer lipids (both on the right side of Fig. 9*B*). Both of these helices became straight gradually such that the entire syntaxin-1 sequences spanning the SNARE motif, jxt linker and TM region formed continuous helices (Fig. 9*C-F*). Such straightening of the helices likely contributes to resolution of the stalk-like structures to form the fusion pore, as the neighboring lipids generally have parallel orientations to the TM regions and therefore re-orient from their initial perpendicular orientation with respect to the flat bilayer to their parallel orientation in the fusion pore (e.g. compare Fig. 7*A* with Fig. 8*B* and 8*F*). At the same time, the re-orientation of the lipids from the distal leaflets of the flat bilayer also brings their polar head groups to the middle to form the fusion pore. The notion that helix continuity from the jxt linker into the TM region of syntaxin-1 plays a role in completing rather than initiating synaptic vesicle fusion is supported by the observation that insertion of a short helix-breaking sequence between the jxt linker and TM region decreases the charge of Ca^2+^-evoked excitatory postsynaptic currents (EPSCs) much less severely than their amplitude because it slowed down the kinetics of release (43).

It is also worth noting that the four-helix bundles of the four SNARE complexes remained bound to the flat bilayer throughout this simulation (Fig. 3*F-H*), as they did in the fusion2g/t350f2 simulations (Fig. 2*B-F*). This feature can be attributed to the multiple positively charged residues of the four-helix bundle that can simultaneously mediate ionic interactions with negatively charged lipids of the flat bilayer, including PIP_2_ (23), and likely dictates the proper orientation of the four-helix bundle with respect to the two membranes (from the point of view of rotation around its long axis) such that the jxt linkers and TM regions of synaptobrevin and syntaxin-1 are suitably projected toward the membrane-membrane interface to efficiently mediate membrane fusion.

### The SNAREs facilitate stalk expansion and fusion pore formation

The mechanism of SNARE-mediate membrane fusion observed in the pullf4link simulation suggests that the SNAREs not only act to bring the membranes together and initiate bilayer merger but also play active roles in promoting the formation of a stalk-like intermediate and generating the fusion pore. This proposal contrasts with results from CG and all-atom simulations with 14-15 nm vesicles suggesting that, once membrane merger is initiated by encounters between acyl chains of the two bilayers at the polar interface, the lipids by themselves can rapidly progress toward formation of the stalk and the fusion pore (35–39).

To distinguish between these two models, we first performed all-atom MD simulations of a 24 nm vesicle and a flat bilayer (Fig. S4). Although the vesicle is still smaller than typical synaptic vesicles [ca. 40 nm (49)], this geometry resembles better that of synaptic vesicle fusion than fusion between two highly curved, 15 nm vesicles.

The simulation was initiated with the vesicle close to the flat bilayer (Fig. S4*A*). The distance between them increased by about 1 nm during 1ns temperature equilibration to 350 K and 1 ns pressure equilibration to 1 atm (Fig. S4*B*), which can be attributed to repulsion between the two membranes. We then ran a 100 ns production simulation at 350 K while applying a pulling force between the acyl chain of one flat bilayer lipid and the acyl chain of a vesicle lipid to promote their encounter at the interface and potentially initiate bilayer fusion. Instead, this pulling force caused the splaying of the vesicle lipid such that one of its acyl chains remained in the vesicle and the other was in the flat bilayer, and the two bilayers came into close contact (Fig. S4*C,E*). To investigate whether such lipid splaying together with the bilayer-bilayer contact might lead to fusion, we extended this simulation for 900 ns more but without the pulling force between the acyl chains (referred to as snfreet350). The splayed lipid remained in this configuration for most of the simulation but was fully in the vesicle at the end (Fig. S4*E-G*). There was no initiation of fusion but, intriguingly, the flat bilayer curved to adapt to the vesicle shape such that one entire side of the flat bilayer made contact with the vesicle (Fig. S4*D,F,G*). This extended interface resembles those observed for SNARE-mediated liposome fusion reactions by cryo-EM (26) and the extended interface that we observed in our first MD simulation with four SNAREs bridging a flat bilayer and a vesicle [Fig. 2J of ref. (23)] but is larger because there are no SNAREs in the periphery. This phenomenon may be driven by entropy, as water molecules become immobilized at membrane interfaces (55) (see discussion).

The snfreet350 simulation suggests that lipid splaying is not sufficient to initiate bilayer merger, which is consistent with the observation that the SNARE-mediated splaying of one lipid in the pullf4link simulation also failed to start fusion (Fig. 5). Instead, fusion was initiated by encounters between acyl chains of the flat bilayer and the vesicle promoted by the SNAREs (Fig. 6). Hence, we asked whether the hydrophobic core induced by the SNAREs that initiated fusion in that simulation might be able to progress unabated toward formation of stalk-like structures, generation of a fusion pore and full fusion without the help of SNARE proteins. For this purpose, we took the 475 ns frame of the pullf4link simulation, in which a substantial hydrophobic core had already formed at the interface, removed the SNARE complexes, replaced their TM regions with lipids and used the resulting system as starting point (Fig. S5*A,B*) for equilibration and production simulation for 900 ns at 350 K (referred to as nscpull4l simulation). Early in the simulation, the flat bilayer curved downwards in the orientation of Fig. S5*C*, away from the vesicle, and the contact between the two bilayers was expanded (Fig. S5*D*). There was some transfer of lipids from the flat bilayer to the vesicle, but no clear expansion of the hydrophobic core was observed by 300 ns of simulation (Fig. S5*D*). Indeed, there was no expansion of the core even at 900 ns, when the size of the core appeared to be smaller and the flat bilayer had wrapped around the vesicle (Fig. S5*E,F*) as we observed for the snfreet350 simulation (Fig. S4*D,G*). Although we cannot rule out the possibility that the hydrophobic core might have expanded at longer time scales, particularly if the curving of the flat bilayer to wrap around the vesicle were avoided, these results reinforce the notion that SNARE proteins facilitate all the steps that lead to fast membrane fusion at a synapse, not only bringing the two membranes together and initiating bilayer merger, but also accelerating formation of stalk-like structures and generating a fusion pore.

## Discussion

The notion that SNARE proteins execute intracellular membrane fusion by forming tight four-helix bundles that bring the two membranes in close proximity has been established for more than two decades, but the molecular mechanism underlying how SNARE complexes induce membrane fusion has remained enigmatic. Elastic continuum models explained the roles of SNAREs and other proteins in membrane fusion in terms of their effects on overall properties such as membrane curvature, elastic moduli and tension (30), and the widespread textbook model of SNARE-mediated membrane fusion envisioned the SNAREs acting as semi-rigid rods that exert mechanical force on the membranes to fuse them as they zipper into SNARE complexes from the N- to the C-terminus (10, 25, 29), which was supported by CG MD simulations (32, 33). Conversely, MD simulations with SNARE-free membranes suggested that fusion is initiated by local encounters of lipid acyl chains at the interface, which may be facilitated by proteins, and membrane fusion can then ensue rapidly without the need of proteins (35–39). Our MD simulations need to be interpreted with caution because of the caveats discussed below but, together with existing experimental data, they suggest a mechanism of how the neuronal SNAREs induce membrane fusion that has a clear, sound physicochemical basis and emphasizes local events at atomic level over general membrane features. In this model, assembly of the SNARE four-helix bundle brings the membranes into close proximity, which is aided at the last stages by the tendency of water to be excluded from the interface. The key event that initiates membrane fusion occurs when the jxt linkers zipper and, together with adjacent hydrophobic residues from the TM regions, they catalyze acyl chain fluctuations out of the bilayers to favor hydrophobic encounters at the polar interface. The resulting small hydrophobic core gradually expands into stalk-like structures and develops into a fusion pore, aided by the TM regions and without a clear intermediate. Our results also suggest that polyunsaturated lipids facilitate the high speed of membrane fusion, which is critical for fast neurotransmission.

Undoubtedly, elastic continuum theory has been very helpful to understand how membranes can fuse (30), and the stalk mechanism has been a guiding light since it was first formulated (56), even though our simulations now suggest that the nature of the stalk at atomic level and its standing as an intermediate in the fusion pathway (in the sense of a defined local energy minimum) may need to be revised (see below). CG MD simulations have also been very insightful to understand membrane fusion but present important caveats, particularly because they cannot simulate substantial conformational changes in proteins [reviewed in (34, 57)]. Conversely, all-atom MD simulations allow the study of structural changes in proteins but are computationally very intensive, limiting the length of time that can be simulated for systems of millions of atoms such as those studied here. Thus, we computed only one simulation in the 1-2 μs time scale for each of the configurations presented; ideally, multiple replicas should be computed, some of them at longer time scales, and additional configurations of the trans-SNARE complexes should be studied to draw firm conclusions. Moreover, the reliability of the simulations depends on the extent to which the force field can reproduce the properties of these systems, and our most important results were obtained at 350 K to help overcome energy barriers; hence, they should be verified with simulations at 310 K that will require much longer times. Nevertheless, it is important to note that the fundamental physicochemical properties of the system are not expected to be substantially altered by the high temperature used apart from the possibility of protein unfolding, which we did not observe. On the contrary, two of the syntaxin-1 molecules became completely helical during the simulation at 350 K that led to SNARE-mediated flat bilayer-vesicle fusion. Importantly, key features of the mechanism of fusion that we observed make a lot of sense from a physicochemical point of view and correlate with extensive experimental data available on neurotransmitter release and liposome fusion (see below).

### Initiation of membrane fusion

Overwhelming evidence shows that SNAREs cannot be viewed as preformed fully helical rods that span the SNARE motifs, jxt linkers and TM regions, and that zipper from N- to C-termini. Thus, isolated SNARE motifs are unstructured (13, 58, 59) and assembly of the SNARE four-helix bundle is templated by Munc18-1 when it binds to synaptobrevin and syntaxin-1, bringing the N-termini of the SNARE motifs together while their C-termini are distant from each other (16, 17) and are flexibly linked to their respective TM regions (60). Moreover, as pointed out above, continuous synaptobrevin and syntaxin-1 helices would be energetically unfavorable in a trans-configuration because of the geometrical reasons. Consistent with these predictions, our attempts to generate trans-SNARE complexes through restrained MD simulations led to loss of helical conformation at the linkers or formation of kinks at the linker helices [(23), Fig. 1*C,D*, 3*A*). Importantly, formation of a continuous helix is clearly not required for the function of synaptobrevin and, actually, flexibility in its TM region is important for release (41, 42, 53, 54) (see below for more details).

In principle, zippering of the four-helix bundle formed by the synapobrevin, syntaxin-1 and SNAP-25 SNARE motifs could be viewed as a mechanical process that overcomes the repulsion between the two membranes and helps to remove the hydration layers between them, events that are commonly assumed to impose substantial energy barriers to membrane fusion [e.g. (30, 31)]. However, the energetic cost of bringing two membranes into contact does not appear to be high, as electrostatic repulsion is very weak at distances beyond 3 nm, cations can shield the charges of lipid head groups and, once two membranes are close, exclusion of water molecules between them actually seems to be favorable from an energetic point of view. The later notion is suggested by the extended membrane-membrane interfaces induced by SNARE complexes in our MD simulations (23), which have been observed experimentally (26), and the way flat bilayers wrap closely around vesicles when held together by lipid bridges (Fig. S4, S5). Moreover, our previous MD simulations suggested that two bilayers could be readily brought into contact even by SNARE complexes in which the C-terminus of the four-helix bundle was not fully assembled (23). It seems likely that water exclusion from the space between two membranes is driven by gains in entropy, as all-atom MD simulations showed that water molecules become ordered at membrane interfaces (55). Mechanical force exerted four-helix bundle assembly may still be important to help push away the abundant membrane proteins present in both membranes from the interface, but our MD simulations suggest that the most important role for the very high stability of the SNARE four-helix bundle is to provide a stable framework to nucleate zippering of the jxt linkers.

Indeed, the most compelling insight from our all-atom MD simulations is that linker zippering constitutes a key event to initiate membrane fusion because it draws the TM regions toward the polar bilayer-bilayer interface and the hydrophobic TM side chains, together with the methylene groups of lysine side chains from the jxt linkers, catalyze the movement of the hydrophobic acyl chains from the lipids to the polar interface (Fig. 5, 6). When acyl chains from the two bilayers come into contact, they form a small hydrophobic nucleus that initiates bilayer merger. Without linker zippering, these encounters are very unlikely, as shown by the fact that we did not observe initiation of membrane merger in MD simulations where SNARE complexes brought two bilayers into contact but the linkers did not zipper [totaling over 15 μs between those described here, those reported in (23), and others that will be described elsewhere]. Note also that the extended contact interfaces between liposomes induced by the SNAREs that were observed by cryo-EM fuse in seconds/minutes (26), showing that high energy barriers hinder linker zippering and spontaneous acyl chain encounters independent of linker zippering under the conditions of these experiments. We did observe acyl chain encounters and merger of the flat bilayer with the vesicle far from the SNARE complex-bridged interface in the t350f2 simulation (Fig. 2) because lipids are not well packed at the edge of the flat bilayer. Similarly, the observation of acyl chain encounters and fusion between closely apposed 14-15 nm vesicles in CG and all-atom MD simulations (35–39) likely arose because of poor lipid packing in these highly curved membranes. The fact that we did not observe such events between 24 nm vesicles and flat bilayers brought together by SNARE complexes without linker zippering indicates that the lipids are sufficiently well packed in both bilayers to make acyl chain encounters at the polar interface very unlikely. Thus, the relatively strong curvature of the 24 nm vesicle, compared to that of synaptic vesicles [ca. 40 nm diameter (49)], does not appear to be a determinant factor for the fusion with the flat bilayer that we observed in the pullf4link simulation.

### Formation of stalk-like structures and progression to a fusion pore

Ample evidence supports the notion that a central step in membrane fusion is the formation of a stalk intermediate in which the proximal leaflets of two bilayers have merged while the distal leaflets have not (30). In our pullf4link simulation, the small hydrophobic nucleus formed by lipid acyl chain encounters at the polar interface between the vesicle and the flat bilayer (Fig. 6*D*) gradually expanded, first at one side of the interface and later toward the middle, forming structures that resemble the postulated stalk intermediate (Fig. 7). However, the interior of these interfacial structures was never fully hydrophobic (Fig. 7*D-F*), as predicted for a stalk, and kept evolving gradually as lipid head groups started to move to the center of the interface and eventually a fusion pore was formed (Fig. 8). Substantial disorder was observed in the lipids at the interface such that some could not be clearly assigned to a leaflet. Such disorder likely was facilitated by the interspersed TM regions of the SNAREs, particularly those of synaptobrevin that were flexibly linked to the jxt linkers throughout the simulation. The exposed polar amide groups of the flexible TM residues likely facilitated the movement of polar head groups to the center of the interface to form the fusion pore. Conversely, the TM regions of two syntaxin-1 molecules formed continuous helices with the jxt linkers and the SNARE motifs relatively early in the simulation, when these helices were bent due to the geometry of the system (Fig. 9*B-D*). Straightening of these helices is expected to be energetically favorable and hence is likely to drag the nearby lipids from their initial orientations approximately perpendicular to the flat bilayer to their parallel orientations in the fusion pore. These observations suggest that, while the TM regions of synaptobrevin and syntaxin-1 may both influence all the events that lead to membrane fusion, the former may play a more preponderant role in facilitating dynamic motions that help the lipids to re-orient during these events, whereas the syntaxin-1 TM region may be more important for ordering the lipids toward formation of the fusion pore structure.

This mechanism does not contradict the stalk hypothesis, as the structures that lead to fusion pore formation do resemble stalks, but they refine the common view of the stalk in three ways. First, there was more lipid disorder than commonly depicted in cartoons of the stalk, which is not surprising because lipid packing is not optimal and is natural that lipids keep moving in search for relatively stable orientations. Second the interior of the stalk-like structures was not purely hydrophobic as expected for protein-free stalks because there are exposed amide groups that interact with lipid head groups. Third, there was no clear intermediate as the system progressed gradually to fusion pore formation. These observations suggest that SNARE-mediated membrane fusion does not involve a unique stalk intermediate corresponding to a well-defined local energy minimum but rather proceeds through a ‘continuum’ of structures that lead to fusion pore formation and might be viewed collectively as a dynamic stalk.

CG and all-atom MD simulations suggested that the defining moment for membrane fusion is the splaying of lipids between the two bilayers and/or encounters of acyl chains from the two bilayers at the interface such as those that we observe, and fusion ensues unabated and quickly (less than 1 μs in most of the simulations) without the help of proteins (35–39). However, these results may also have arisen from poor packing of the highly curved 14-15 nm vesicles used in these studies, as we did not observed fusion of the flat bilayer to the 24 nm vesicle when they were bridged by a splayed lipid or by a hydrophobic nucleus involving multiple contacts between acyl chains from the two bilayers (Fig. S4, S5). While it is plausible that fusion might have been observed at much longer time scales, these results support the notion emerging from our pullf4link simulation that the SNARE TMs play an active role in accelerating all the steps of membrane fusion from initial bilayer-bilayer contact to fusion pore formation, as was suggested previously by CG MD simulations (61).

### Experimental support and implications for the mechanisms underlying neurotransmitter release and membrane fusion

It is satisfying that the mechanism of SNARE-mediated membrane fusion suggested by our MD simulations correlates with and explains multiple lines of available experimental evidence. Particularly important are the findings that insertion of a helix-breaking proline-proline sequence between the SNARE motif and the jxt linker of synaptobrevin (85PP) strongly impaired liposome fusion (42) and insertion of another helix-breaking sequence (glycine-serine-glycine) between the SNARE motif and jxt linker of syntaxin-1 (259GSG) practically abolished neurotransmitter release (43, 44). These results showed that jxt linker zippering plays a crucial role for liposome fusion in vitro and neurotransmitter release in neurons, in correlation with the key importance of linker zippering for the fusion mechanism that we observed in our simulations. In contrast, a PP insertion between the jxt linker and the TM region region of synaptrobrevin (93PP) had no or small effects on liposome fusion in vitro (42) and Ca^2+^-triggered exocytosis in chromaffin cells (41), showing that zippering into the synaptobrevin TM region is not functionally important and consistent with the observation that TM residues close to the jxt linker remained flexible throughout our pullf4link simulation. In fact, the functional defects caused by mutations in the helix-breaking glycine residue of the synaptobrevin TM region (G100) or mutations that stabilize helical conformation showed that conformational flexibility in the synaptobrevin TM region is crucial for efficient Ca^2+^-triggered exocytosis (53, 54). Conversely, a GSG insertion between the jxt linker and the TM region of syntaxin-1 (265GSG) did decrease the amplitude of EPSCs in neurons dramatically although it had little effect on the EPSC charge because it slowed down the kinetics of release (43). These results can be naturally explained by the observation that extension of the jxt linker helix into the TM of syntaxin-1 was not important to initiate membrane fusion but such helical extension likely facilitated formation of the fusion pore in our pullf4link simulation. Furthermore, the strong impairment of Ca^2+^-triggered exocytosis in chromaffin cells caused by addition of two charged residues (lysines or glutamates) at the C-terminus of synaptobrevin led to the conclusion that the C-terminus needs to be pulled inside the vesicle membrane during exocytosis (62), and such pulling is a direct consequence of linker zippering as observed in the pullf4link simulation.

The 259GSG and 265GSG insertions in syntaxin-1 lead to 75% and 60% reductions in release induced by hypertonic sucrose, which is used to measure the readily-releasable pool of synaptic vesicles (43). These results indicate that linker zippering also facilitates release in the slower time scales of these experiments but is not as crucial as it is for fast Ca^2+^-evoked release, and that extension of the jxt helix into the TM region also helps but is not essential for sucrose-induced release. Spontaneous neurotransmitter release was not substantially altered by the 259GSG insertion but was enhanced three-fold by the 265GSG insertion (43, 44), indicating that spontaneous release does not require linker zippering but somehow depends on coupling between the jxt linker and the TM region of syntaxin-1.

Overall, the available data suggest that zippering of the synaptobrevin and syntaxin-1 linkers is particularly crucial for Ca^2+^-triggered neurotransmitter release because it helps to trigger membrane fusion in microseconds, as observed in our MD simulations. Linker zippering is expected to be hindered by energy barriers, as it pulls the TM regions and involves conformational changes in the linkers, which should be unstructured or form kinked helices in the primed state [(23), Fig. 1*D*, 3*D*]. Moreover, linker zippering likely requires remodeling of interactions between the lipids and the abundant basic and aromatic residues present in the jxt linkers (23, 63), which is supported by the effects of mutations in the linkers on release (64, 65). The substantial folding energy associated with linker zippering (47) likely helps to overcome these energy barriers, but it is tempting to speculate that linker zippering is also facilitated during Ca^2+^-triggering of release by other components of the neurotransmitter release machinery (e.g. synaptotagmin-1) through an as yet unknown mechanism.

Linker zippering also facilitates fusion at longer time scales such as those involved in sucrose induced release (43, 44) and bulk liposome fusion assays (42). Moreover, reconstitution experiments with the vacuolar fusion machinery have shown that the jxt linkers of the corresponding SNAREs also play a key role in liposome fusion (45) and the jxt linker of the exocytotic syntaxin homologue in yeast is important for secretion (46). Hence, it seems likely that the overall notion that linker zippering promotes hydrophobic encounters of the lipid acyl chains of two bilayers at the interface to initiate bilayer merger constitutes a general principle underlying intracellular membrane fusion. The earlier finding that surfactant-associated protein B also promoted such encounters to initiate fusion of 14 nm vesicles in CG MD simulations (39) suggests that these principles likely apply also to other forms membrane fusion, including viral induced fusion. Our MD simulations do not support models predicting a proteinaceous fusion pore [e.g. (66, 67)] but do indicate that the SNARE TM regions play active roles during membrane fusion and hence provide frameworks to reinterpret the mutagenesis data that suggested that these regions line the fusion pore in these studies. Note also that the importance of the TM regions for fusion has been controversial, as replacing them with lipid anchors allowed neurotransmitter release or SNARE-mediated liposome fusion in some studies but not in others (44, 68–70). The mechanism of SNARE-mediated membrane fusion uncovered by our MD simulations suggests that the ability of lipid anchors to support fusion may depend on the way they are attached to the SNAREs and whether they attach hydrophobic moieties close enough to the jxt linkers such that they can facilitate lipid acyl chain encounters at the interface.

An additional insight provided by our pullf4link simulation is that the acyl chain encounters at the interface primarily involved lipids with polyunsaturated acyl chains. Such lipids are highly abundant in synaptic vesicles (49) and brain membranes in general (50), are known to be important for brain health (71) and they were shown to enhance liposome fusion in vitro (72), which likely arises because of mismatch between the length and conformational preferences of the saturated acyl chain commonly present in the same lipid together with the polyunsaturated chain. Our results suggest that polyunsatured lipids may be important for the fast speed of Ca^2+^-evoked neurotransmitter release.

Clearly, further research will be necessary to test the overall model of SNARE-mediated membrane fusion and other ideas presented here. Our results provide a vivid illustration of the power of all-atom MD simulations to make further progress in this field and to elucidate complex biological problems.

## Methods

### High performance computing

All-atom MD simulations were performed using Gromacs (51, 52) with the CHARMM36 force field (73). Most high performance computing, including all production MD simulations were carried out on Frontera at TACC. System setup, including solvation, ion addition, minimizations and equilibration steps were performed at the BioHPC supercomputing facility of UT Southwestern. System visualization and manual manipulation were performed with Pymol (Schrödinger, LLC).

### System setup and MD simulations

The methodology used to set up systems, equilibrate them and run production MD simulations was analogous to that described previously (23). The vesicle generated previously (23) was adapted for each system by moving lipids manually to accommodate different positions of the SNARE TM regions or their absence. A flat square bilayer of 30.5 nm x 30.5 nm was built at the CHARMM-GUI website (48) (https://charmm-gui.org/) for the fusion2g system and was adapted for all other systems (also moving lipids manually to accommodate SNARE TM regions) except for the snfreet350 system, which used a smaller bilayer adapted from the qscv simulation of ref. (23). The lipid compositions of the original vesicle and flat bilayers are described in Table 1, and those of each system were only slightly altered by adding or removing a few lipids as needed. The number of atoms, box dimensions and simulation temperatures of each system are also listed in Table 1. All systems were solvated with explicit water molecules (TIP3P model), adding potassium and chloride ions as needed to reach a concentration of 145 mM and make the system neutral.

All systems were energy minimized, heated to the desired temperature over the course of a 1 ns MD simulation in the NVT ensemble with 1 fs steps, and equilibrated to 1 atm for 1 ns in the NPT ensemble using isotropic Parrinello-Rahman pressure coupling and 2 fs steps (74). NPT production MD simulations were performed for the times indicated in Table 1 using 2 fs steps, isotropic Parrinello-Rahman pressure coupling and a 1.1 nm cutoff for non-bonding interactions. Nose-Hoover temperature coupling (75) was used separately for three groups: i) protein atoms; ii) lipid atoms; and ii) water and KCL. Periodic boundary conditions were imposed with Particle Mesh Ewald (PME) (76) summation for long-range electrostatics. The speeds of the production simulations ran on Frontera at TACC were typically about 28 ns/day for a typical system of about 5.2 million atoms using 40 nodes.

## Acknowledgments

We thank Carsten Kutzner and Helmut Grubmüller for advice on how to optimize the parameters of our MD simulations. Most of the work presented in this paper was performed through Pathways and Leadership Computing Resource allocations on Frontera at the Texas Advanced Computing Center of The University of Texas at Austin (URL: http://www.tacc.utexas.edu) (projects MCB20033 and IBN23002). This research also used computational resources provided by the BioHPC supercomputing facility located in the Lyda Hill Department of Bioinformatics, UT Southwestern Medical Center, TX (URL: https://portal.biohpc.swmed.edu). This work was supported by grant I-1304 from the Welch Foundation (to JR) and by NIH Research Project Award R35 NS097333 (to JR).

## Supplementary Figures

**Figure S1.**
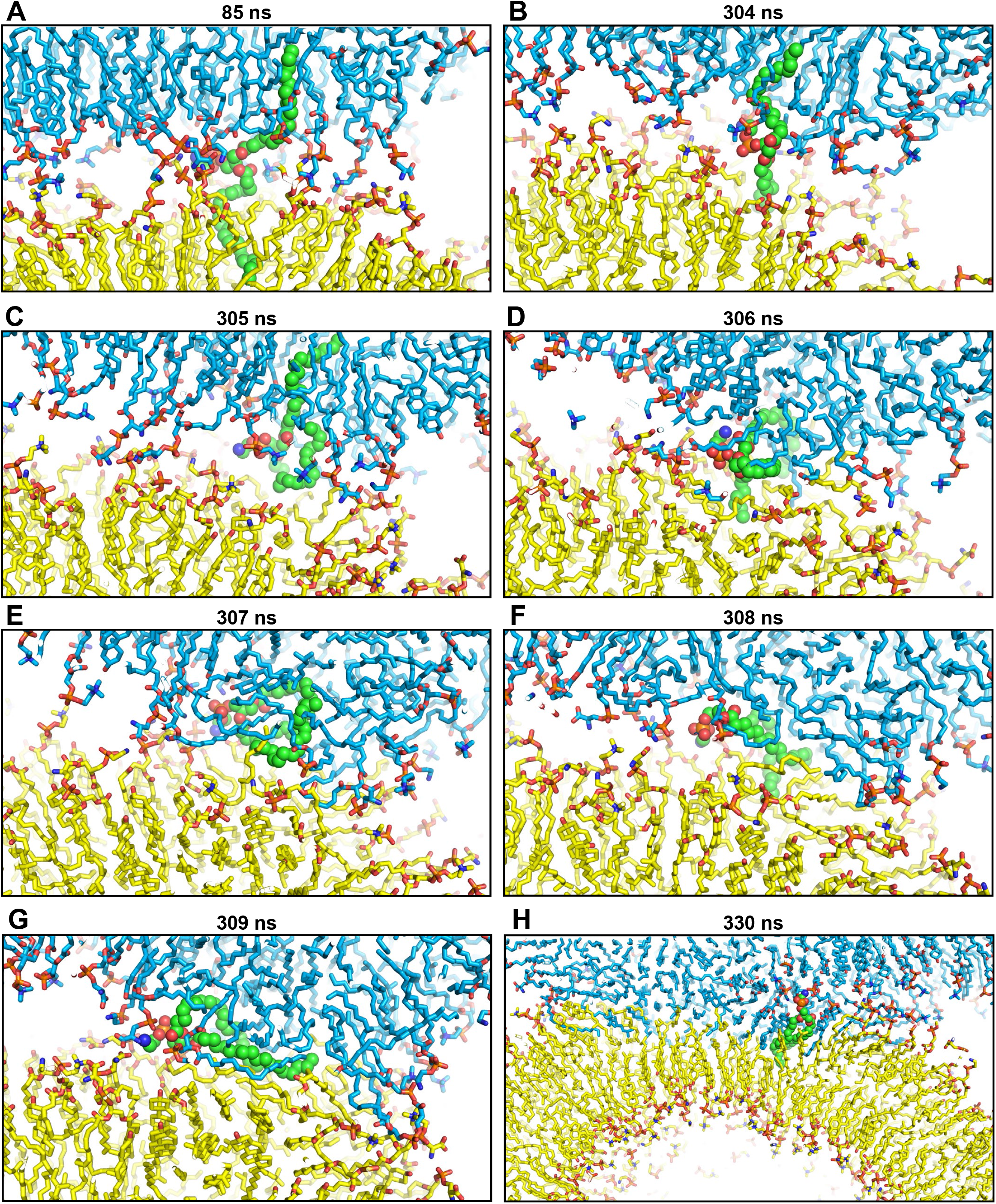
Snapshots taken at the indicated time points of the t350f2 simulation showing the splaying of an SDPE molecule at the interface between the edge of the flat bilayer and the vesicle, and the merger of the bilayers. Note that the SDPE molecule remained splayed for over 200 ns without the membrane merging. Membrane merger occurred when fluctuations of the SDPE molecule led to hydrophobic encounters at the polar interface. Lipids and proteins are shown as stick models, and SNARE complexes are in addition represented by ribbon diagrams. The color-code is the same as in Fig. 2. The SDPE molecule is represented by spheres with nitrogen atoms in dark blue, oxygens in red, phosphorus in orange and carbon in green.

**Figure S2.**
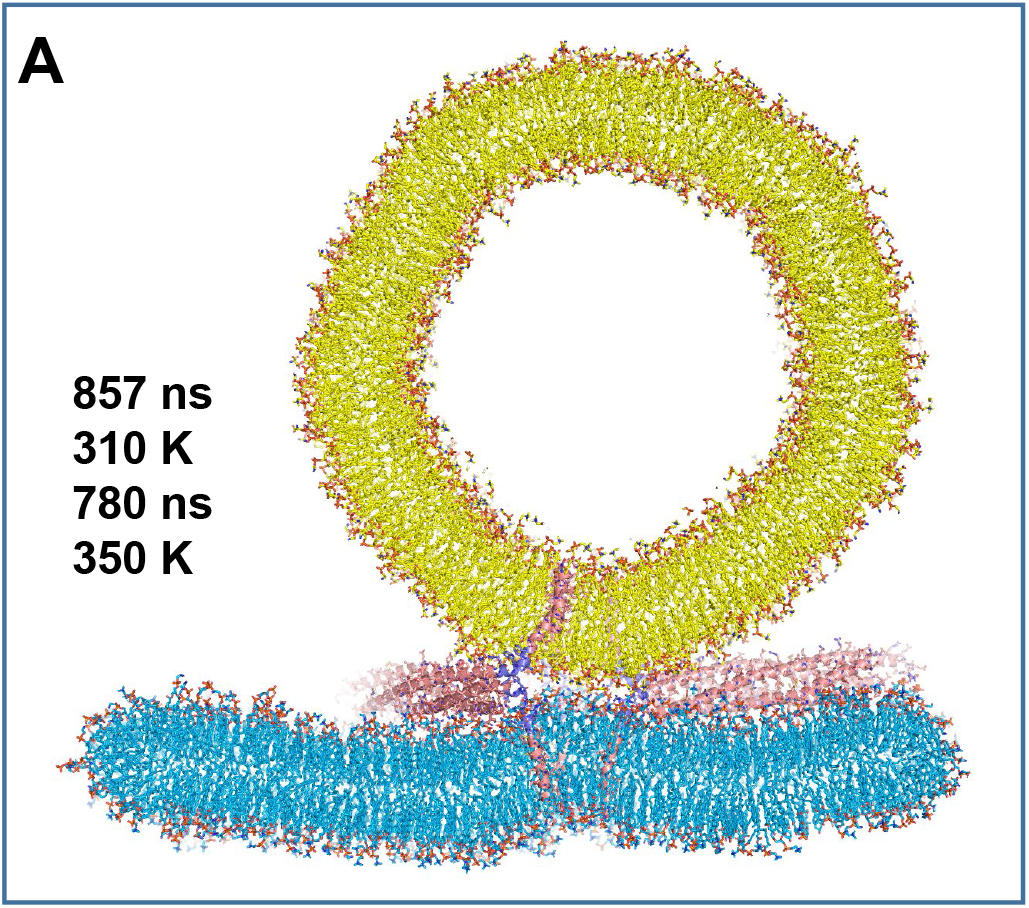
Slice showing the t350f2e system after 780 ns of simulation at 350 K. Lipids and proteins are shown as stick models, and SNARE complexes are in addition represented by ribbon diagrams. The color-code is the same as in Fig. 2.

**Figure S3.**
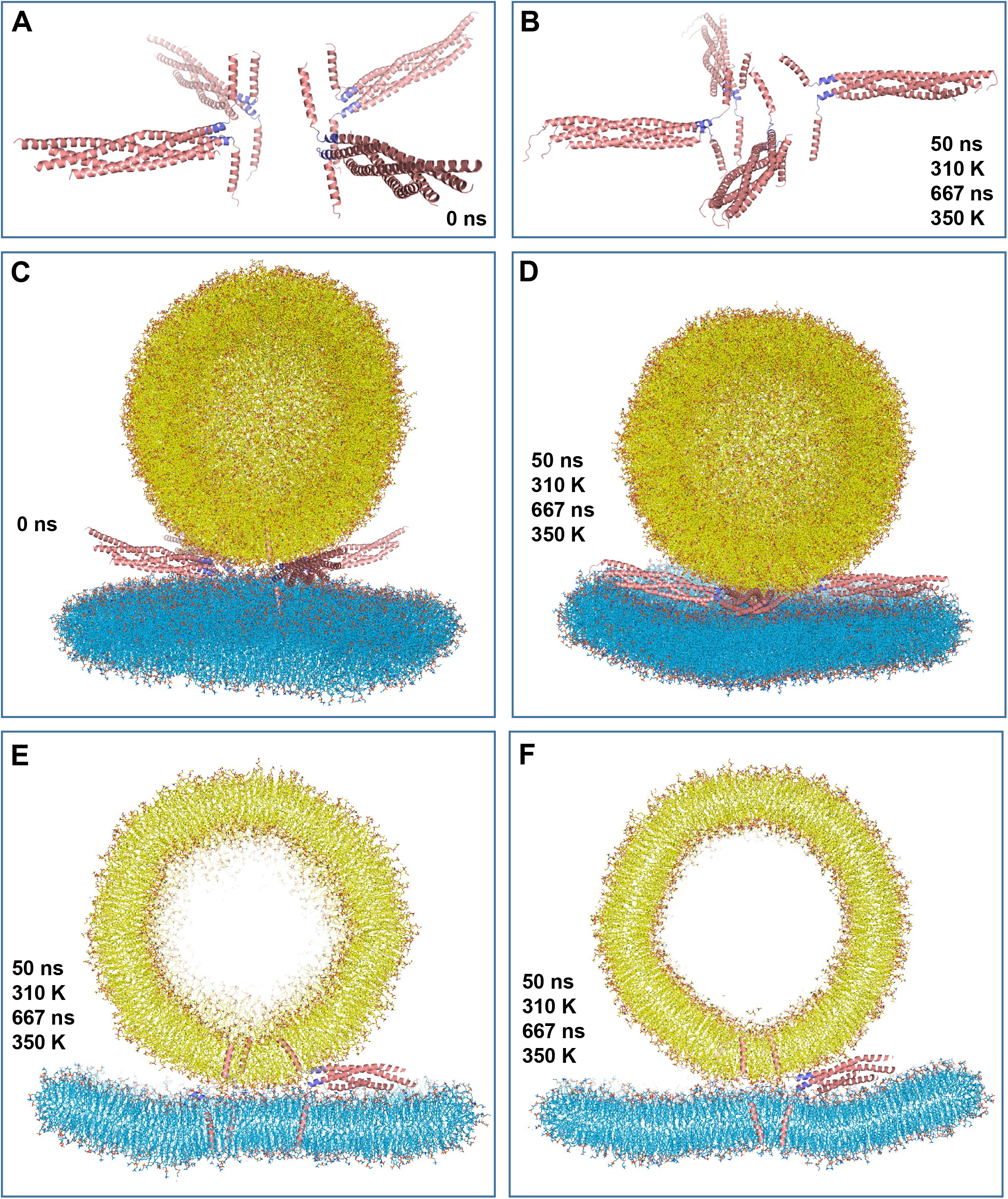
MD simulation of four trans-SNARE complexes bridging a vesicle and a flat bilayer, with the linkers zippered but no pulling force to keep them zippered (t350f4link simulation). (*A*) Ribbon diagram showing initial configuration of the trans-SNARE complexes used for this simulation, which resulted from restrained MD simulations to zipper the linkers starting with the complexes shown in Fig. 3A. (*B*) Ribbon diagram of the trans-SNARE complexes at the end of the simulation. (*C-D*) Full views of the system at the indicate time points of the t350f4link simulation. (*E-F*) Slices of the system at the end of the simulation taken from different angles to show how the flat bilayer was somewhat buckled in one direction but not another. In (*C-F*), Lipids and proteins are shown as stick models, and SNARE complexes are in addition represented by ribbon diagrams. The color-code is the same as in Fig. 2.

**Figure S4.**
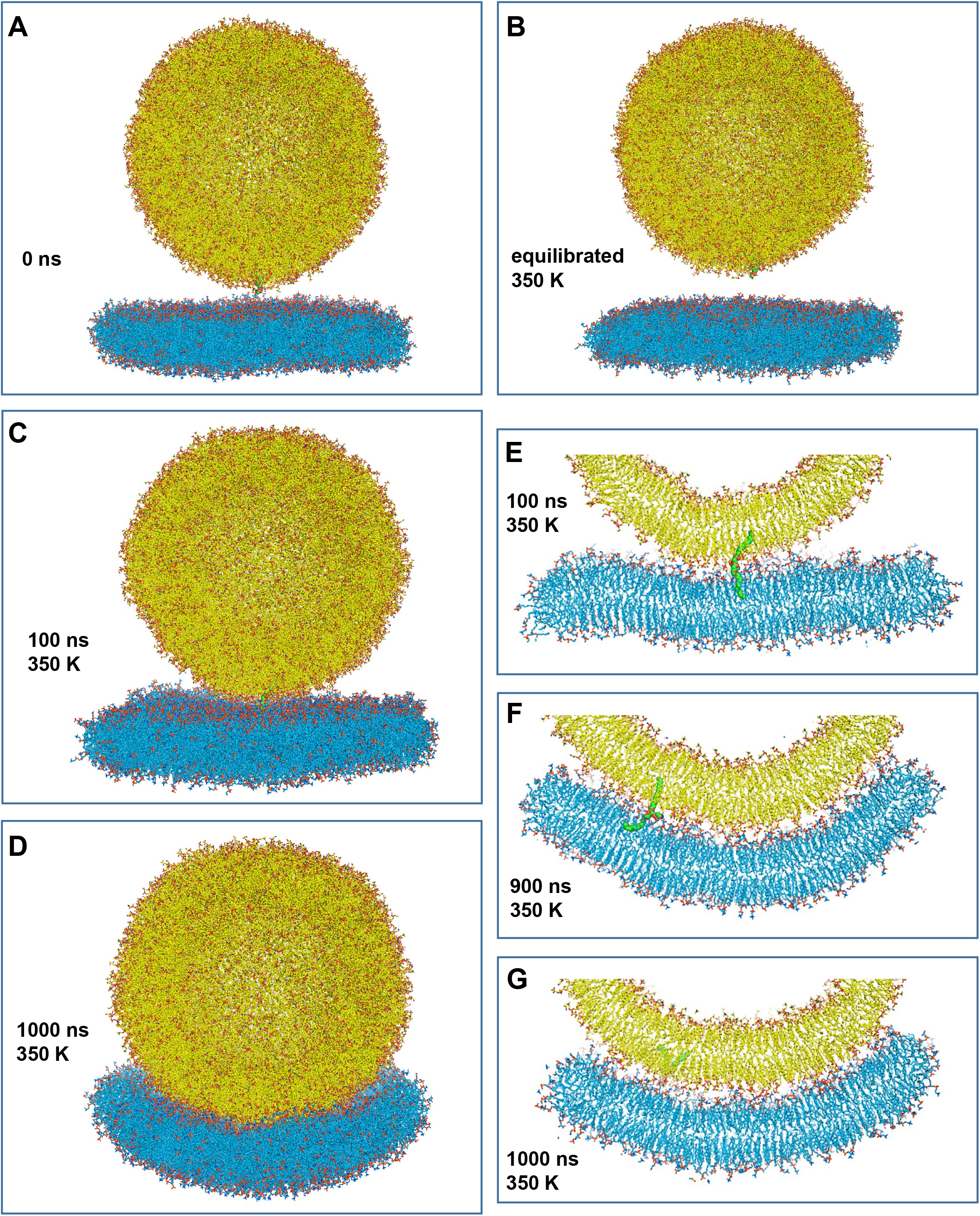
Simulation of a flat bilayer and a vesicle bridged by a splayed lipid. (*A-D*) Full views of the initial system (*A*), the system after temperature equilibration for 1 ns and pressure equilibration for 1 ns (*B*), and after 100 ns (*C*) or 1000 ns (*D*) of production simulation at 350 K. (*E-F*) Slices taken of the system taken at the indicated time points. Lipids and proteins are shown as stick models, and SNARE complexes are in addition represented by ribbon diagrams. The color-code is the same as in Fig. 2. The splayed lipid is represented by spheres with nitrogen atoms in dark blue, oxygens in red, phosphorus in orange and carbon in green.

**Figure S5.**
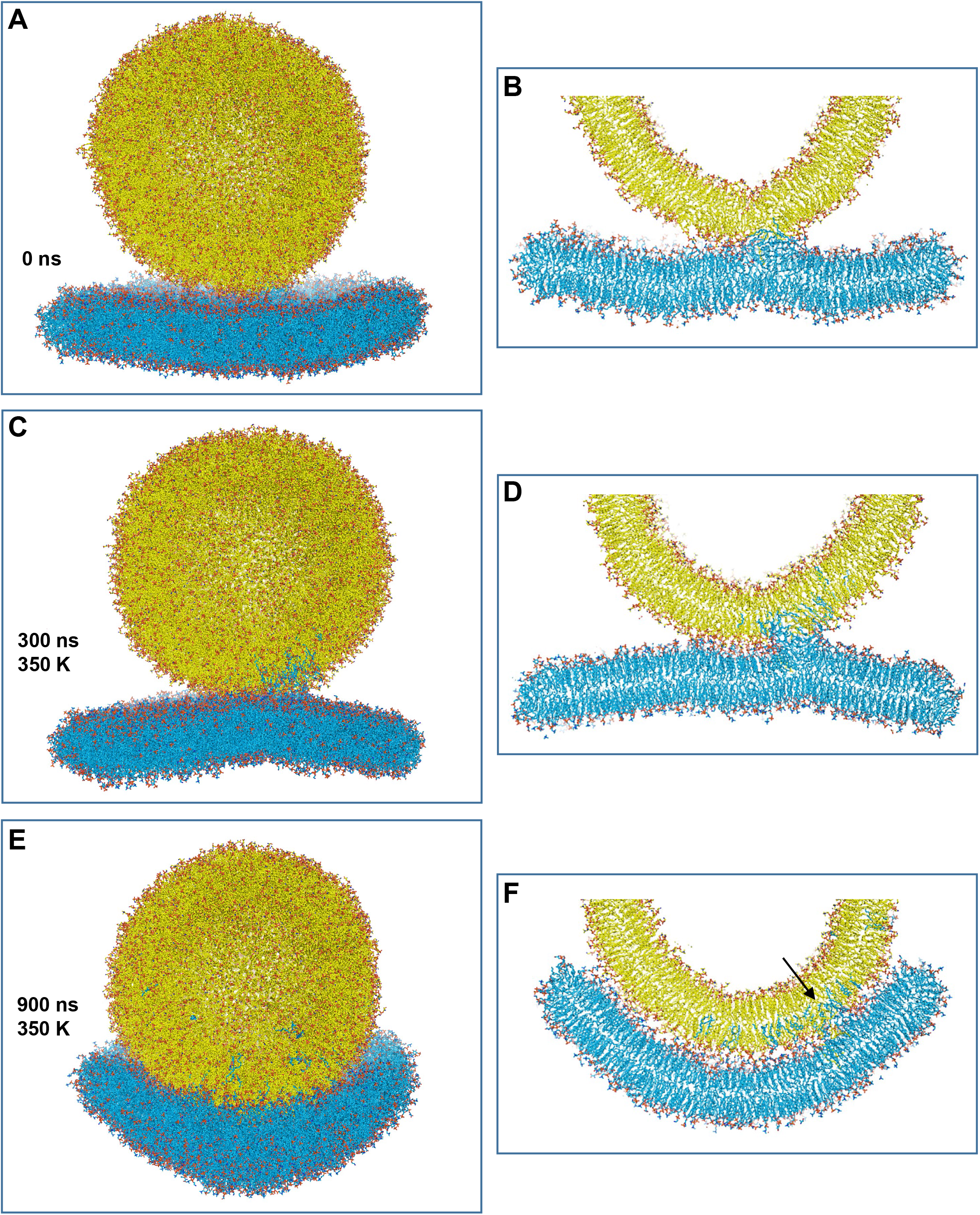
Simulation of a flat bilayer bridged to a vesicle by a few lipids that form a small hydrophobic core at the interface. (*A*) Initial configuration of the system, which was generated from the 475 ns frame of the pullf4link simulation by removing the SNAREs and replacing their TM regions with lipids. (*C,E*) Full view of the system after 300 ns (*C*) and 900 ns (*E*) of production simulation at 350 K. (*B,D,F*) Slices of the system at the stages shown in panels (*A,C,E*), respectively. The slice at 900 ns shows how the entire flat bilayer wraps around the vesicle and some of its lipids have been transferred to the vesicle. The arrow of panel (*F*) points at the hydrophobic core remaining at the polar interface.

